# Human intestinal organoid-derived PDGFRα+ mesenchymal stroma empowers LGR4+ epithelial stem cells

**DOI:** 10.1101/2023.08.16.553630

**Authors:** Chen JunLong, Shinichiro Horiuchi, So Kuramochi, Tomoyuki Kawasaki, Hayato Kawasumi, Saeko Akiyama, Tomoki Arai, Kenichi Morinaga, Tohru Kimura, Tohru Kiyono, Hidenori Akutsu, Seiichi Ishida, Akihiro Umezawa

**Affiliations:** Center for Regenerative Medicine, National Center for Child Health and Development Research Institute, Tokyo, Japan; Department of Advanced Pediatric Medicine, Tohoku University School of Medicine, Sendai, Japan; Division of Pharmacology, National Institute of Health Sciences, Kawasaki, Japan; 1st Section, 1st Development Department, Food and Healthcare Business Development Unit, Business Development Division, Research & Business Development Center, Dai Nippon Printing Co., Ltd., Tokyo, Japan; Laboratory of Stem Cell Biology, Department of BioSciences, Kitasato University School of Science, Kanagawa, Japan; Project for Prevention of HPV-Related Cancer, Exploratory Oncology Research and Clinical Trial Center, National Cancer Center, Chiba, Japan; Graduate School of Engineering, Sojo University

**Keywords:** mesenchyme, intestinal stem cells, pluripotent stem cells, organoids, trophocytes, GREM1, regeneration, adherent culture, epithelium-mesenchyme co-culture, pharmacokinetics

## Abstract

The columnar epithelial cells comprising the intestinal tract, stomach, and uterus can be cultured in vitro as organoids or in adherent culture. However, the proliferation of these columnar epithelial cells in adherent culture is limited. Likewise, human pluripotent stem cell (hPSC)-derived intestinal epithelial cells do not show extensive or clonal propagation in vitro. In this study, we induced proliferation of hPSC-derived small intestinal epithelium for a longer time by utilizing mesenchymal stromal cells derived from self-organized intestinal organoids as feeders. The proliferating cells exhibited columnar form, microvilli and glycocalyx formation, and cell polarity, as well as expression of drug-metabolizing enzymes and transporters. It is noteworthy that small intestinal epithelial stem cells cannot be cultured in adherent culture alone, and the stromal cells cannot be replaced by other feeders. Organoid-derived mesenchymal stromal cells resemble the trophocytes essential for maintaining small intestinal epithelial stem cells, and play a crucial role in adherent culture. The high proliferative expansion, productivity, and functionality of hPSC-derived small intestinal epithelial stem cells could have potential applications in pharmacokinetic and toxicity studies and regenerative medicine.

## INTRODUCTION

The small intestine constitutes the largest organ that facilitates the assimilation of nutrients and conveys it to the bloodstream throughout the body. Intestinal epithelial cells boast a plethora of transporters and metabolic enzymes, such as CYP3A4, a crucial enzyme in the arena of drug discovery, and play a pivotal role in the absorption and metabolic processing of drugs.^1,2^ As such, it is imperative to assess the gastrointestinal tract’s capacity for absorbing and metabolizing drugs during the drug discovery phase. The mainstream approach to drug discovery primarily involves the utilization of cells derived from experimental animals, Caco2 cells, and primary human cells, yet there remain concerns regarding possible disparities in results owing to differences in species and gene expression between the epithelial cell models and human intestine.^3,4^

Pluripotent stem cells, such as embryonic stem cells and induced pluripotent stem cells, are pluripotent and immortal. Clinical applications include disease models and regenerative medicine, while drug discovery applications include pharmacokinetic studies and organ-on-chip models. The advantage of pluripotent stem cells is that they can be used indefinitely by differentiating them into desired cell types in vitro, but complex induction protocols and the lack of passaging of differentiated cells are major challenges. Of particular concern are the lot-to-lot differences that occur when differentiating from pluripotent stem cells and when differentiated cells are repeatedly passaged and cryopreserved. During the process of proliferation in adherent culture, epithelial cells lose their morphology, barrier function, and metabolic capacity and must maintain their function as three-dimensional structures called "organoids".^5–8^ These organoids are artificially fabricated to resemble living organs and exhibit highly functional tissue bodies.

Two major types of organoids are currently reported due to differences in fabrication methods. One is reconstructed organoids, in which parenchymal cells, stromal cells, and vascular endothelial cells are differentiated from pluripotent stem cells, mixed, and embedded in an extracellular matrix to create a three-dimensional organoid.^5–8^ The other is self-organized organoids (self-organization).^9^ The generation process of the self-organized organoids is based on the principles of embryonic development, in which different cell types organize in a highly coordinated manner to form complex structures such as organs. Organoids can reproduce many of the characteristics and functions of the corresponding organs, such as the production of specific hormones or specific metabolic functions. Self-organizing organoids are excellent models for studying organ development and disease because they better reproduce the natural cellular organization and structure of organs. They can also spontaneously generate a greater variety of cell types, which may lead to a higher degree of complexity and functionality.

Intestinal epithelial stem and progenitor cells are influenced by subepithelial signals such as Wnt and BMPi (GREM1).^10–13^ Recent advances in high-resolution microscopy and single-cell RNA sequencing have revealed that subepithelial myofibroblasts,^14–16^ i.e. telocytes and trophocytes, are a source of mesenchymal trophic factors.^17,18^ These myofibroblasts in the mucosal layer have been extensively studied with respect to niche function. Co-culture with myofibroblasts contributes to the proliferation of intestinal organoids via Wnt, Rspo, and BMPi.^12,14^ These myofibroblasts are Vimentin**^+^** α-SMA**^+^** Desmin^-^, and trophocytes, one of the myofibroblasts are CD81^+^ PDGFRA^low^.^18^

Mesenchymal stromal cells play a crucial role in the maintenance of epithelium in many tissues and organs. Mesenchymal stromal cells facilitate the long-term culture of human hepatocytes through Wnt3a and R-spondin 1.^19^ This study highlights the utility of mesenchymal cells derived from intestinal organoids to support human LGR4-positive intestinal stem cells.

## RESULTS

### Intestinal epithelial cells can be isolated from organoids

The intestinal epithelial cells were generated from hPSC-derived self-organized intestinal organoids^9^. The organoids exhibited balloon-like structures 1 cm in diameter and were maintained as floating foam (Figures 1A and 1B). Intestinal organoids had well-developed stromal cell layers and exhibited peristaltic-like movements, indicating highly functional small intestinal organoids (Figure 1C). Intestinal organoids had a complex tissue structure, i.e. both epithelial and stromal cell layers (Figures 1D and 1E). To collect intestinal epithelial cells, organoids were cut and processed into single sheets (Fragment, Figure 1F). Then the mesenchyme surface was adhered to a dish coated with laminin (Figures 1G and 1H). On the first day, a minimum amount of medium was used to prevent intestinal organoid fragments from floating. The medium was returned to normal volume after confirming the adhesion of the intestinal organoids to the dish.

**Figure 1.**
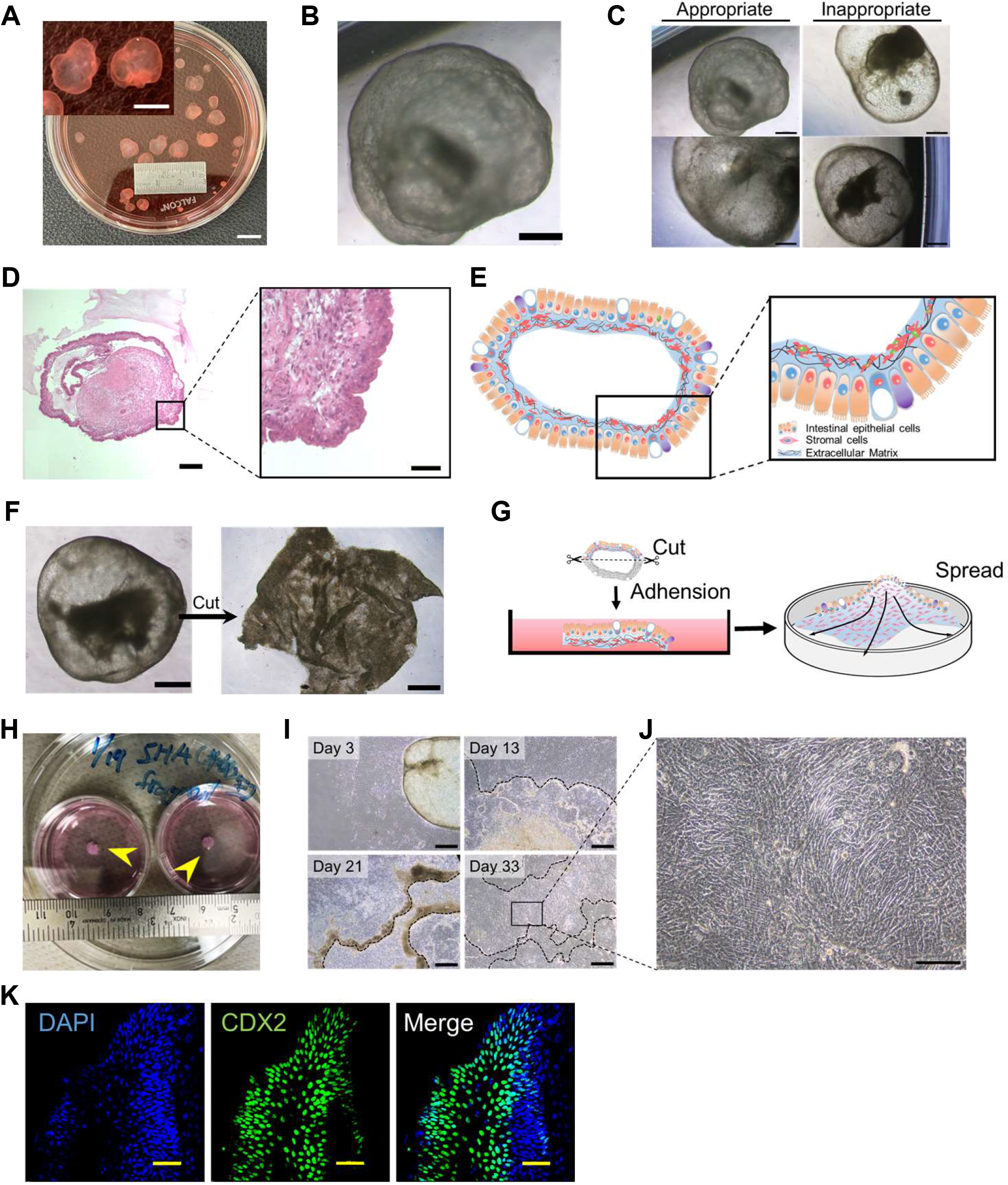
Isolation of intestinal epithelial cells from small intestinal organoids. (A) Macroscopic view of intestinal organoids. Bar: 10 mm. Inset: High-power view. Bar: 5 mm. (B) Phase-contrast micrograph of small intestinal organoids with submucosa. Scale bar: 500 µm. (C) Phase-contrast photomicrographs of appropriate (left) and inappropriate (right) ’intestinal organoids’. Organ-derived small intestinal epithelium and mucosal stroma can be obtained from the appropriate organoids that have thick cell layers and low light transmission. Bar: 500 μm. (D) Histology of small intestinal organoids with submucosa. HE stains. Scale bars: 200 (left), 50 (right) µm. (E) Schematic of small intestinal organoids with submucosa. (F) Phase-contrast micrographs of an organoid cut open and unfolded with a scissor. Scale bars: 500 μm. (G) Three-dimensional small intestinal organoids were bisected to expose the submucosa and attached to the dish. Cells were spread on the dish. (H) Macroscopic view of the dissected organoids that are attached to the dishes. (I) Phase-contrast micrograph of a bisected organoid (black dotted lines) on 3, 13, 21, and 33 days after adhesion. Scale bars: 500 µm. (J) Phase-contrast photomicrographs of proliferating intestinal epithelial cells on the mucosal stroma. Scale bar: 100 µm. (K) Immunocytochemistry of adherent organoids with an antibody to CDX2, an intestinal epithelial marker. Nuclei were stained with 4’,6-diamidino-2-phenylindole dihydrochloride solution (DAPI). Scale bars: 100 µm.

Mesenchymal stromal cells with a typical spindle shape started to migrate and proliferate from the adherent organoids within 7 days after the adhesion and covered the entire dish within 2 weeks (Figure 1I). The intestinal epithelial cells proliferated in the form of colonies over 80% of the dish in 30 days (Figure 1I). The cells resembled fetal-derived intestinal epithelial cells in morphology (Figure 1J).^20^ Interestingly, colonies of intestinal epithelial cells cultured in organoid production medium were maintained without cell proliferation for more than 60 days (data not shown). In addition, intestinal organoid-derived epithelial cells expressed CDX2 (Figure 1K).

We employed a medium containing TGF-β inhibitor, Cholera toxin, Nicotinamide, EGF, Noggin, Wnt3a and R-spondin1 that support the proliferative expansion of a diverse array of human epithelial cells.^19,21–24^ This study demonstrates that it is also efficacious for intestinal epithelial cells.

### Mesenchymal stromal cells support organoid-derived epithelial cells upon passage

We passaged the epithelial cells onto a mouse embryonic fibroblast (MEF) feeder. However, the epithelial cells failed to maintain their cell morphology and lost CDX2 expression at the center of the colonies (Figures 2A, 2B and S1A). CDX2 expression was only observed at the periphery of the colonies, indicating that MEFs as a feeder are inappropriate for supporting intestinal epithelial cells. We then isolated mesenchymal stromal cells from intestinal organoids and examined their attributes. We defined "intestinal mesenchymal stromal cells" and "intestinal epithelial cells" isolated from intestinal organoids as "LONG" and "RYU", respectively, as code names. The mesenchymal stromal cells were positive for Vimentin and α-SMA but negative for Desmin (Figure 2C), showing a property of subepithelial myofibroblasts.^14–16^ The mesenchymal stromal cells exhibited robust expression of trophocyte markers PDGFRA, CD81, and GREM1, while devoid of CD34 (Figures 2D and 2E).^18,25^ Mesenchymal stromal cells efficiently proliferated (Figure 2F) but ceased to divide at passage 15, probably due to replicative senescence (Figure S1B). Mesenchymal stromal cells highly expressed RSPO3, BMP inhibitor (GREM1, FST), and WNT5A (non-canonical WNT) (Figures 2G and 2H), which are key trophic factors in maintaining epithelial stemness.^12,18,26^

**Figure 2.**
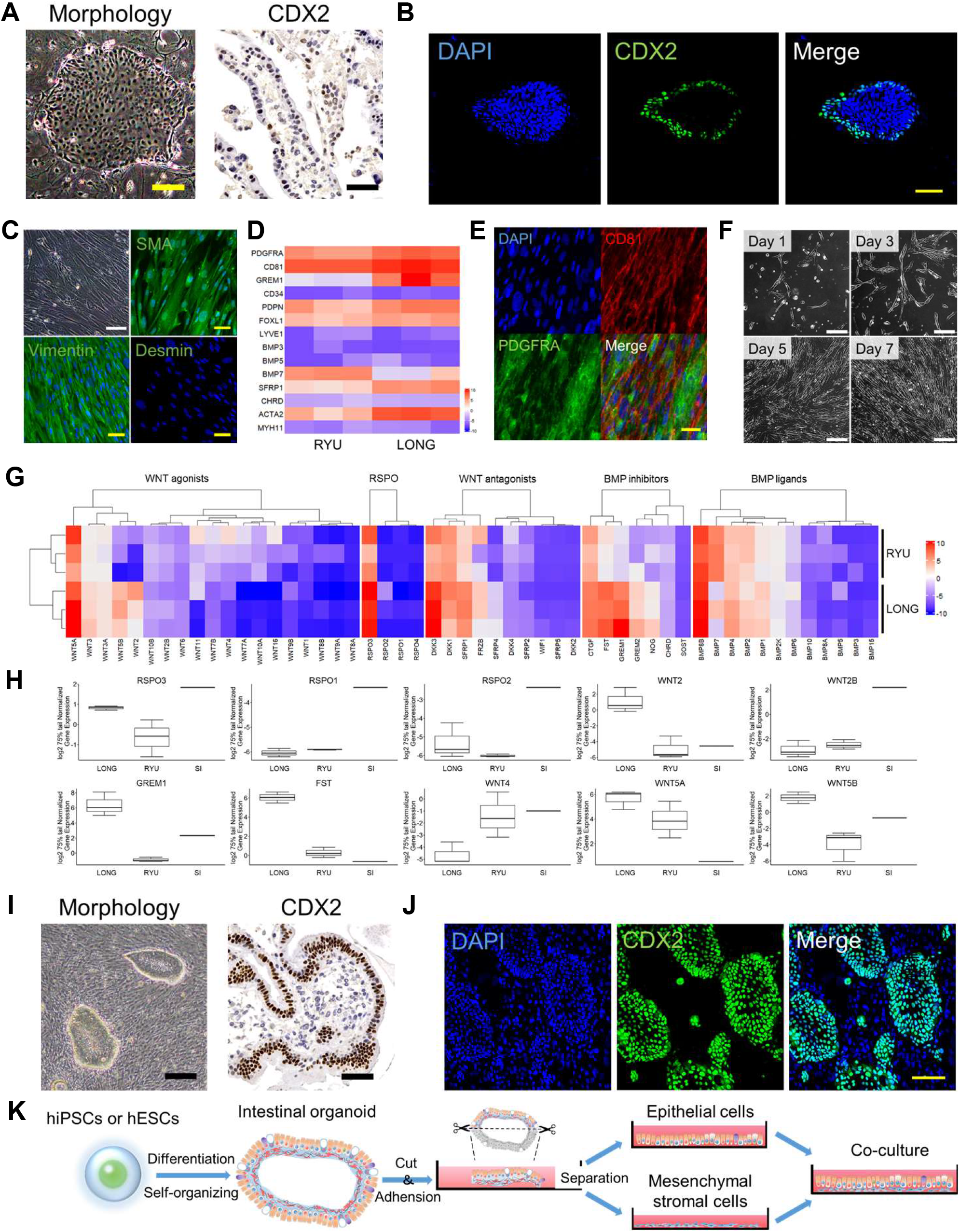
Serial culture of epithelial cells derived from small intestinal organoids. (A) Phase-contrast micrograph of intestinal epithelial cells at passage 2 on mouse embryonic fibroblasts (MEF) (left) and immunohistochemistry of intestinal epithelial cells with an antibody to CDX2 (right). Immunohistochemistry was performed on cells in iPGell. Scale bars: 200 (left) and 50 (right) µm. (B) Immunocytochemistry of intestinal epithelial cells at passage 2 on MEF with an antibody to CDX2. Nuclei were stained with DAPI. Scale bar: 100 µm. (C) Phase-contrast micrograph of and immunohistochemistry of mesenchymal stromal cells. Mesenchymal stromal cells exhibit spindle-shaped morphology. Mesenchymal stromal cells were positive for Vimentin and α-SMA, but negative for Desmin. Nuclei were stained with DAPI. Scale bars: 100 (white) and 50 (yellow) µm. (D) Expression heat map of gene expression in intestinal epithelial cells and mesenchymal stromal cells. (E) Immunocytochemistry of mesenchymal stromal cells. Mesenchymal stromal cells were positive for trophocyte markers PDGFRA and CD81. Nuclei were stained with DAPI. Scale bar: 50 µm. (F) Phase-contrast micrograph mesenchymal stromal cells. Days after a passage are shown. Scale bars: 200 µm. (G) Trophic factor gene expression of intestinal epithelial cells and mesenchymal stromal cells. (H) Boxplot of representative Rspo, WNT, and BMP inhibitor families. Each expression level was calculated from the results of independent (biological) triplicate experiments (intestinal epithelial cells and mesenchymal stromal cells) and single experiments (normal adult intestine tissue). Boxplots are expressed as mean ± SD. (I) Phase-contrast micrograph and immunohistochemistry of intestinal epithelial cells on organoid-derived mesenchymal stromal cells with an antibody to CDX2. Immunohistochemistry was performed on cells in iPGell. Scale bars: 200 (left) and 50 (right) µm. (J) Immunocytochemistry of intestinal epithelial cells on organoid-derived mesenchymal stromal cells with an antibody to CDX2. Nuclei were stained with DAPI. Scale bars: 100 µm. (K) Schematic of successful maintenance and culture of intestinal epithelial cells on organoid-derived mesenchymal stromal cells. Epithelial cells and mesenchymal stromal cells were derived from the same organoids.

We then investigated whether mesenchymal stromal cells support intestinal epithelial cell passaging. The intestinal epithelial cells maintained a typical columnar epithelial cell morphology with clear cell boundaries and formed colonies with CDX2-positive cells at 100% (Figures 2I and 2J). Mesenchymal stromal cells as a feeder significantly rescued the decrease in CDX2 positive cells (Figures 2I, 2J and S1C). It is also noteworthy that mesenchymal stromal cells increased the adhesion of intestinal epithelial cells. Indeed, intestinal epithelial cells showed a low rate of adhesion to the dish in the absence of mesenchymal stromal cells, and the cells that did adhere showed a rapid loss of epithelial characteristics (Figure S2). Therefore, mesenchymal stromal cells play an essential role in maintaining characteristics and passaging intestinal epithelial cells, which previously could not be cultured in adherent culture. Likewise, mesenchymal stromal cells also supported intestinal spherical organoids using the standard established method (Figures S3A and S3B).^7^ The establishment of mesenchymal stromal cells and intestinal epithelial cells is schematically illustrated (Figures 2K and S4A).

### Proliferation and genomic stability of epithelial cells on mesenchymal stromal cells

We investigated the proliferative capability of human embryonic stem cells (ESC) and induced pluripotent stem cells (iPSC)-derived intestinal epithelial cells from 3D organoid technology (Figures 3A and S4B). A large number of mesenchymal stromal cells were cryopreserved for further use of feeder cells. Intestinal epithelial cells, after propagation on mesenchymal stromal cells, can easily be passaged, cryopreserved, and thawed. The intestinal epithelial cells were passaged from one to four dishes every 5-7 days (Figure 3B and Supplemental video). A marker for proliferating cells, Ki67, was detected in intestinal epithelial cells (Figure 3C). Intestinal epithelial cells continued to proliferate until 30 population doublings, i.e., 2^30^ cells from 1 cell, in 120 days and stopped dividing (Figures 3D–3H). The intestinal epithelial cells lacked expression of the stem cell marker LGR5 but highly expressed LGR4, CD24, CD44, ZNF277, and SOX9 (Figures S5A, S5B and S5C). Hence, the intestinal epithelial cells correspond to transient amplifying cells in the crypt base, but not intestinal stem cells.^20,27–29^ Intestinal epithelial cells showed a normal karyotype with high chromosomal stability (Figure 3I). Intestinal epithelial cells and mesenchymal stromal cells formed a typical luminal structure with epithelial cells and stromal cells at the subrenal capsules of immunodeficient (NOD.Cg-PrkdcscidIl2rgtm1Sug/ShiJic) mice.^30^ Moreover, there was no evidence of tumorigenicity for either of the cell types (Figures 3J and S5D).

**Figure 3.**
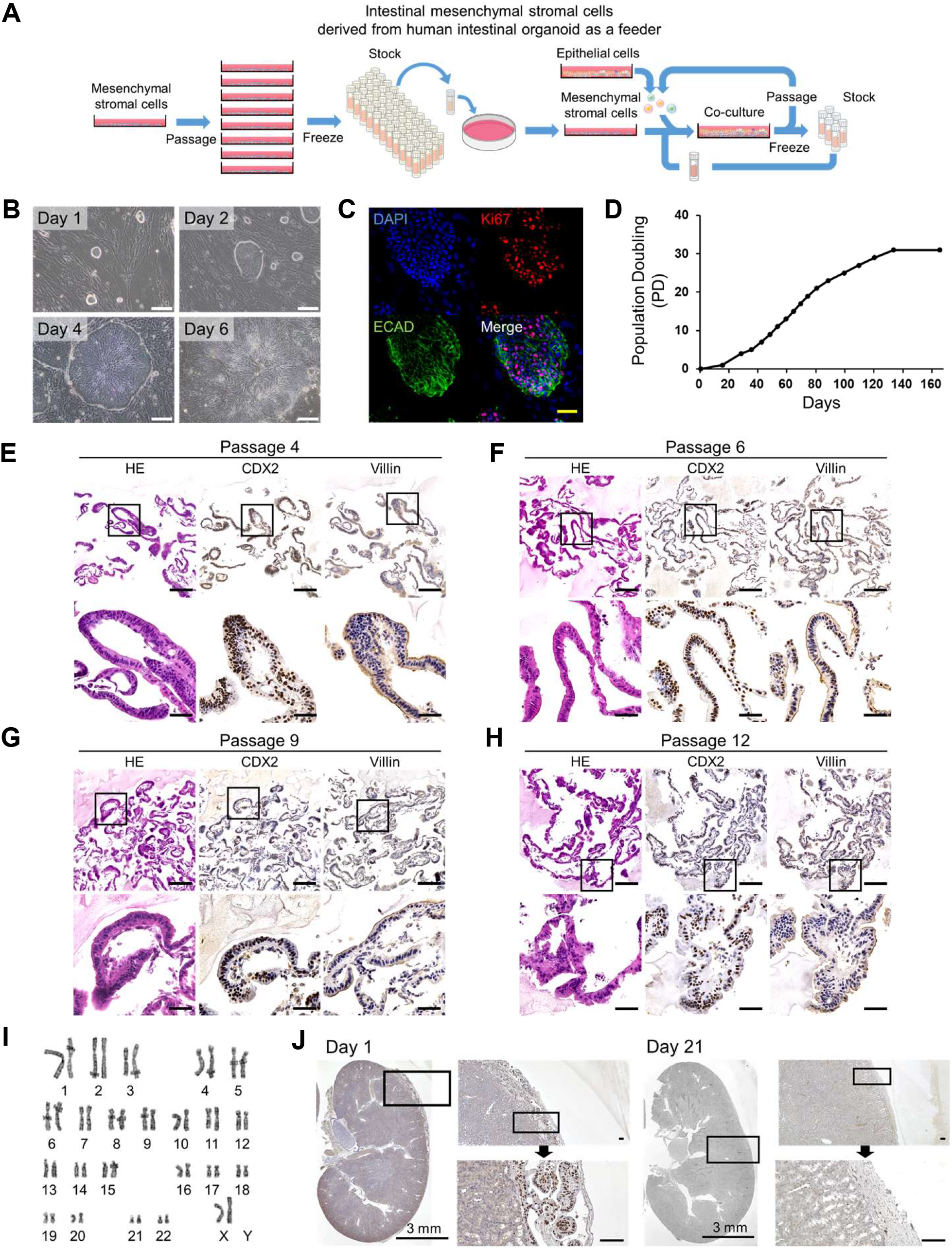
Extensive proliferation of organoid-derived intestinal epithelial cells on a stromal cell feeder layer. (A) Schematic of organoid-derived intestinal epithelial cell culture system. Intestinal epithelial and mesenchymal stromal cells can be cultured, passaged and frozen, allowing for large stocks of cells. Intestinal epithelial and mesenchymal stromal cells are designated as RYU and LONG, respectively, as code names. (B) Phase-contrast micrographs of intestinal epithelial cells growing on mesenchymal stromal cells. The epithelial cells formed colonies and proliferated. Scale bars: 100 µm. (C) Immunocytochemistry of intestinal epithelial cells on a dish with the proliferating cell marker Ki67 and the epithelial cell-specific cadherin (ECAD (E-Cadherin/Cadherin-1/CD324)). Nuclei were stained with DAPI. Scale bar: 50 µm. (D) Growth curve of small intestinal epithelial cells. Each point indicates the timing of passages. The cells were passaged into 4 dishes at each passage. (E) Histology and immunohistochemistry of intestinal epithelial cells at passage 4 in iPGell. Immunohistochemistry was performed with antibodies to CDX2 and Villin. Scale bars: 50 µm. (F) Histology and immunohistochemistry of intestinal epithelial cells at passage 6 in iPGell. Immunohistochemistry was performed with antibodies to CDX2 and Villin. Scale bars: 50 µm. (G) Histology and immunohistochemistry of intestinal epithelial cells at passage 9 in iPGell. Immunohistochemistry was performed with antibodies to CDX2 and Villin. Scale bars: 50 µm. (H) Histology and immunohistochemistry of intestinal epithelial cells at passage 12 in iPGell. Immunohistochemistry was performed with antibodies to CDX2 and Villin. Scale bars: 50 µm. (I) Karyotypic analysis revealed that organoid-derived intestinal epithelial cells at passage 13 have normal karyotypes. (J) Immunohistochemistry of organoid-derived intestinal epithelial cells and mesenchymal stromal cells at the subrenal capsule of the kidney, using human-specific Lamin A/C antibody. Scale bars of zoomed sections represent 100 µm.

### Intestinal epithelial cells exhibit polarization in adherent culture

Immunostaining and scanning electron microscopy analysis revealed that intestinal epithelial cells proliferated on mesenchymal stromal cells (Figures 4A and 4B). The structure of these cells in the adhesion culture resembled the epithelial mucosal layer of the small intestine and intestinal organoid. Histological analysis revealed that the intestinal epithelial cells had a typical columnar shape and formed villus structures (Figure 4C). The epithelial cells were positive for CDX2 and epithelial keratins (AE1/3), and mesenchymal stromal cells were positive for Vimentin. The intestinal epithelial cells expressed Villin and transporters (BCRP, PEPT1, and P-gp) on the apical membrane side, showing cell polarity (Figure 4D). The intestinal epithelial cells produced acid mucus in the apical membrane and intracellular vacuoles (Figure 4E). The vacuoles were positive for MUC2, specific for goblet cells in the small intestine (Figure 4F). Electron microscopy analysis clearly revealed microvilli lined with actin filaments and a glycocalyx on the apical membrane (Figures 4G and 4H). Intestinal epithelial cells are composed of absorptive enterocytes and goblet cells on two or more-layered mesenchymal stromal cells and show cell polarity (Figure 4I).

**Figure 4.**
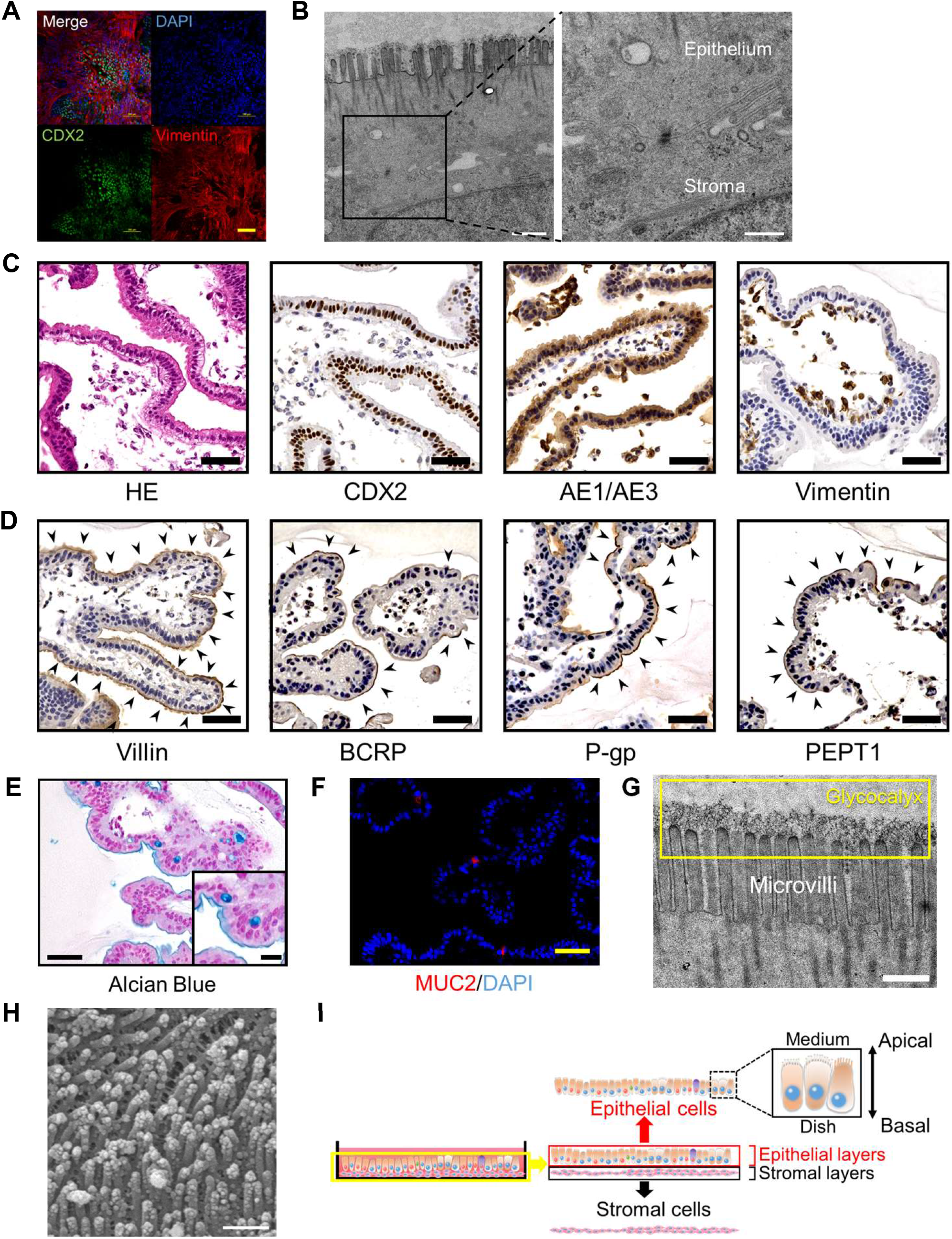
Organoid-derived intestinal epithelial cells have polarity. (A) Immunocytochemistry of intestinal epithelial cells on mesenchymal stromal cells with the antibody to Vimentin and CDX2. Epithelial and mesenchymal stromal cells are positive for CDX2 and Vimentin, respectively. Scale bar: 100 µm. (B) Transmission electron microscopic analysis of intestinal epithelial cells on mesenchymal stromal cells. Adhesion between epithelial and stromal cells was observed. Scale bars: 1000 (left) and 500 nm. (C) Histology and immunohistochemistry of intestinal epithelial cells in iPGell. Immunohistochemistry was performed with antibodies to CDX2, AE1/AE3, and Vimentin. Scale bars: 50 µm. (D) Immunohistochemistry of intestinal epithelial cells in iPGell was performed with antibodies to Villin, BCRP, P-gp, and PEPT1. Strongly stained areas are indicated by black arrows. Scale bars: 50 µm. (E) Alcian Blue stain of intestinal epithelial cells iPGell. Scale bars: 50 and 20 (inset) µm. (F) Immunohistochemistry of intestinal epithelial cells in iPGell with antibodies to MUC2. Scale bar: 50 µm. (G) Transmission electron microscopic analysis of epithelial apical membrane side. Microvilli and glycocalyx (yellow squares) were observed. Scale bar: 200 nm. (H) Scanning electron microscopic analysis of intestinal epithelial cells on mesenchymal stromal cells. Microvilli were observed over the surface of intestinal epithelial cells. Scale bar: 500 nm. (I) Schematic of intestinal epithelial cells on mesenchymal stromal cells. Intestinal epithelial cells are polarized in adherent culture. The apical and basal side of intestinal epithelial cells is in contact with culture medium and mesenchymal stromal cells, respectively.

### The intestinal epithelial cells have barrier function and P-gp ability

The intestinal tract serves as a selectively permeable barrier, which regulates the absorption of both nutrients and xenobiotics. Intestinal epithelial cells’ barrier function and capacity to transport P-glycoprotein were quantified. The intestinal epithelial cells on mesenchymal stromal cells exhibited positivity for ECAD and CDH17, which are elements of adherens junctions. Likewise, the cells were positive for ZO-1, Claudin-2 and Claudin-7, constituents of tight junctions (Figure 5A). Transmission electron microscopy analysis revealed tight junctions and adherens junctions at the lateral membranes of the intestinal epithelial cells (Figure 5B). We then evaluated the barrier function of the intestinal epithelial cells by transepithelial electrical resistance (TEER) measurements and Lucifer yellow (LY) assays (Figures 5C and 5D). The TEER value for the intestinal epithelial cells on mesenchymal stromal cells was 235-311 Ω cm² (Figure 5C). The Lucifer yellow apparent permeability (Papp) value was 0.45-0.98 x 10□□ cm/s for the intestinal epithelial cells on mesenchymal stromal cells (Figure 5D). These results demonstrate that the intestinal epithelial cells possess a high barrier function comparable or superior to Caco-2 cells. Subsequently, we examined the time course of intestinal epithelial cells’ barrier function for 20 days (Figure 5E). The intestinal epithelial cells on mesenchymal stromal cells demonstrated a consistent barrier function (Papp value: 1.5 - 2.6 x 10□□ cm/s) throughout the 20 days.

**Figure 5.**
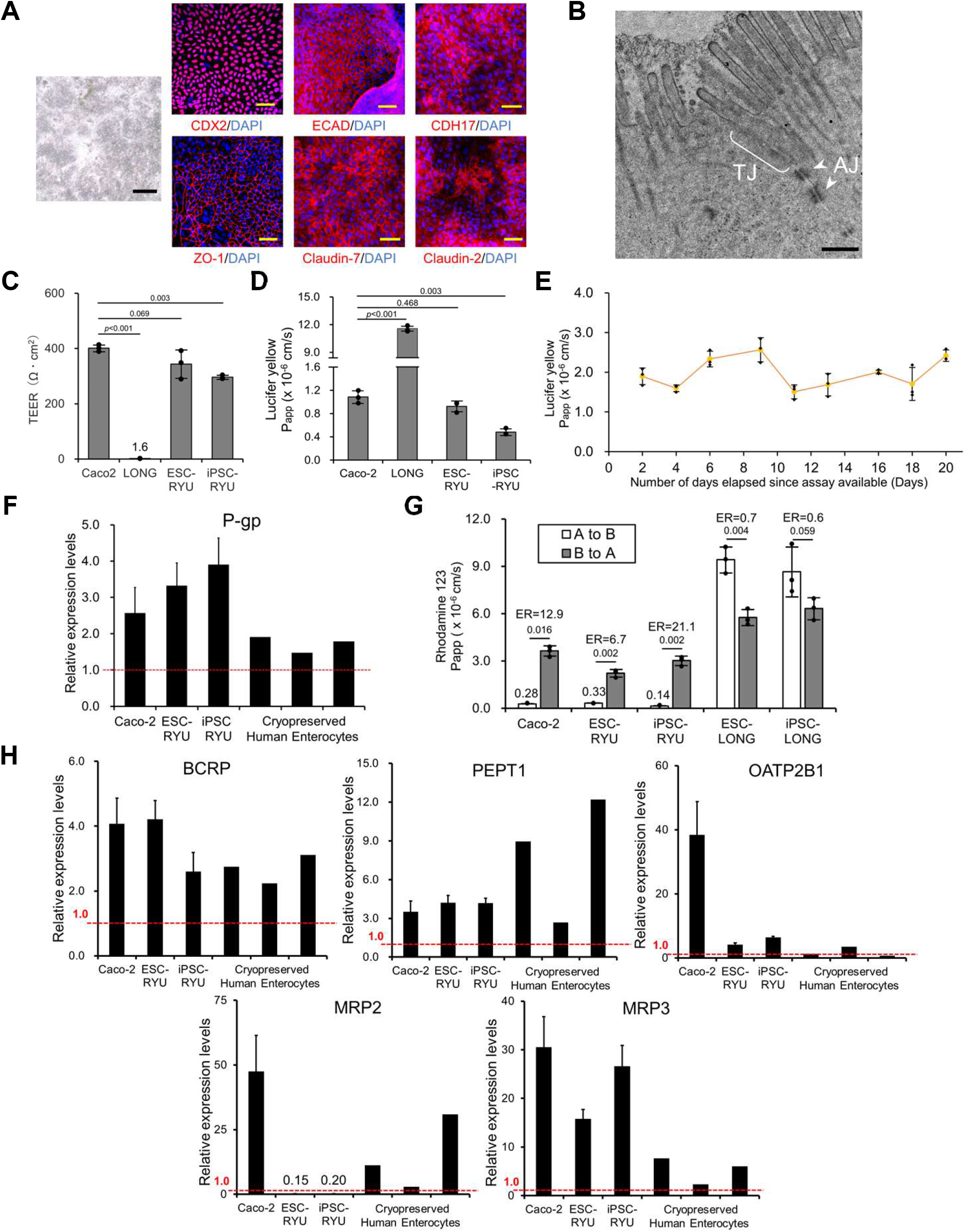
Barrier performance and transport capability of intestinal epithelial cells. (A) Phase-contrast micrograph and immunocytochemistry of epithelial cells on mesenchymal stromal cells. Immunocytochemistry was performed with the antibody to barrier functions (ECAD, CDH17, ZO-1, Claudin-2, Claudin-7) and CDX2. Nuclei were stained with DAPI. Scale bars: 500 (black) and 50 (yellow) µm. (B) Transmission electron microscopic analysis of epithelial lateral membrane. Tight junction (TJ) and adherens junction (AJ, white arrows) were observed. Scale bar: 500 nm. (C) Trans-epithelial electrical resistance measurements of Caco-2, mesenchymal stromal cells (LONG), and intestinal epithelial cells derived from human iPSCs (iPSC-RYU) and ESCs (ESC-RYU) on mesenchymal stromal cells. Mesenchymal stromal cells and Caco-2 cells served as negative and positive controls, respectively. Results are expressed as mean ± SD (n = 3 triplicate biological experiments). Statistical significance was determined using Dunnett’s test compared to Caco-2. (D) Lucifer Yellow permeability test of intestinal epithelial cells derived from human iPSCs (iPSC-RYU) and ESCs (ESC-RYU) on mesenchymal stromal cells. Caco-2 and mesenchymal stromal cells (LONG) were prepared as controls. ESC-RYU: ESC-derived intestinal epithelial cells, iPSC-RYU: iPSC-derived intestinal epithelial cells. Results are expressed as mean ± SD (n = 3 triplicate biological experiments). Statistical significance was determined using Dunnett’s test. (E) Lucifer Yellow permeability test was performed for barrier performance. Barrier performance of epithelial cells on mesenchymal stromal cells. Results are expressed as mean ± SD (n = 3 triplicate biological experiments). (F) P-glycoprotein (P-gp) gene expression by qRT-PCR analysis in intestinal epithelial cells derived from human iPSCs and ESCs on mesenchymal stromal cells in a transwell. Caco-2 cells and cryopreserved human enterocytes served as controls. The expression level of adult small intestine whole tissue was set to 1.0. Results are expressed as mean ± SD (n = 3 triplicate biological experiments). (G) Permeability test with Rhodamine 123. The efflux ratio (ER) for Rhodamine 123 was derived from the Papp values associated with basal-to-apical transport (B to A) and apical-to-basal transport (A to B). Intestinal epithelial cells derived from human iPSCs and ESCs on mesenchymal stromal cells. Mesenchymal stromal cells alone (ESC-LONG and iPSC-LONG) and Caco-2 cells served as negative and positive controls, respectively. Results are expressed as mean ± SD (n = 3 triplicate biological experiments). Statistical significance was determined using Student’s t-test. (H) Expression of the genes for transporters (BCRP, PEPT1, OATP2B1, MRP2, and MRP3). The expression level of adult small intestine whole tissue was set to 1.0. Each expression level was calculated from the results of independent (biological) triplicate experiments. Results are expressed as mean ± SD (n = 3). ES-RYU: human ESC-derived intestinal epithelial cells, iPS-RYU: human iPSC-derived intestinal epithelial cells.

The intestinal epithelial cells and Caco-2 cells had the barrier function. Then, we confirmed the expression levels of intestinal transporters, and the transporter ability was examined. P-glycoprotein (P-gp) plays a crucial role in the intestinal absorption and excretion of drugs.^31^ The intestinal epithelial cells express comparable levels of P-gp than Caco-2 cells and cryopreserved human enterocytes (Figure 5F). We performed a transport assay using Rhodamine123 (P-gp substrate) in the intestinal epithelial cells (Figure 5G). The intestinal epithelial cells on mesenchymal stromal cells exhibited asymmetric permeability of P-gp substrate Rhodamine123, with efflux ratios (ER) 6.7-21.1. Mesenchymal stromal cells exhibited an efflux ratio (ER) ranging from 0.6-0.7, which, in conjunction with the gene expression findings, indicates their non-P-gp activity. These results suggest that the intestinal epithelial cells possess high P-gp activity comparable or superior to Caco-2 cells. The intestinal epithelial cells expressed other transporters, BCRP, PEPT1, OATP2B1, and MRP3, whereas the expression of MRP2 was limited. (Figure 5H). The expression of transporters except MRP2 was comparable to or superior to cryopreserved human enterocytes (IVAL-E1, E2, and E3).

### Cytochrome P450 is induced in the intestinal epithelium by drugs

Among the cytochrome P450 enzymes (CYPs), CYP3A4 is a dominant drug-metabolizing enzyme;^32^ carboxylesterase 2 (CES2) contributes to the pharmacokinetics of oral drugs possessing an ester structure in the epithelial cells of the small intestine.^33,34^ However, the expression levels of CES2 and CYP3A4 in Caco-2 cells are significantly lower than those observed in the human adult small intestine.^35^ We evaluated the CYP gene expression and functions in the intestinal epithelial cells. The intestinal epithelial cells expressed CYP1A1, CYP2B6, CYP2C9, CYP2C19, CYP2D6, and CYP3A4, but not CYP1A2 (Figure 6A). The expression of CYPs was comparable to or superior to Caco-2 cells. The intestinal epithelial cells possessed the same level of CYP3A4 expression as cryopreserved human enterocytes. The intestinal epithelial cells exhibited distinct expression patterns when compared to CYP-expressing hepatocytes (Figure S6A and S6B). Furthermore, the expression of CYPs in intestinal epithelium is induced by drugs.^36,37^ In the current study, we sought to examine the CYP induction of intestinal epithelial cells. The intestinal epithelial cells on mesenchymal stromal cells exhibited induction of CYPs with exposure to Omeprazole(OM), β-Naphthoflavone (BNF), Phenobarbital (PB), Rifampicin (Rif), vitamin D (VD3), and Dexamethasone (Dex) through a wide range of nuclear receptors, i.e., PXR, VDR, GR, and AHR (Figure 6B and S6C).^38,39^ Furthermore, the intestinal epithelial cells exhibited deficient expression of the nuclear receptor CAR, similar to that observed in intestinal organoids.^40^ The intestinal epithelial cells on mesenchymal stromal cells exhibited inhibition of CYP3A4 with exposure to Ketoconazole (Ket) (Figure S6D). The intestinal epithelial cells were positive for CYP3A4 and CES2 (Figure 6C).

**Figure 6.**
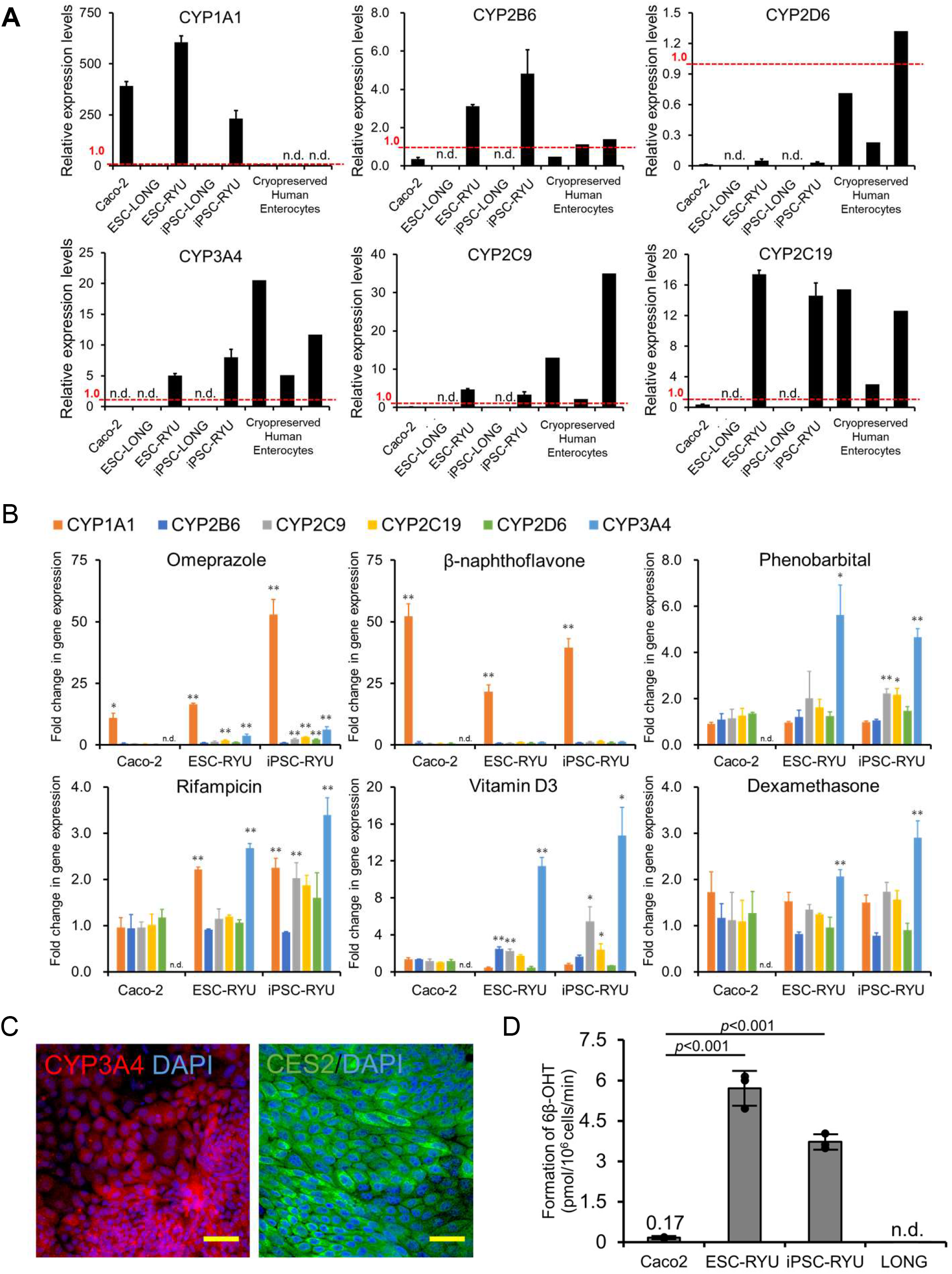
Induction of the genes for cytochrome P450 in organoid-derived epithelial cells. (A) Expression of the genes for cytochrome P450 (CYP1A1, CYP2B6, CYP2D6, CYP3A4, CYP2C9, and CYP2C19). The expression level of adult small intestine whole tissue was set to 1.0. Each expression level was calculated from the results of independent (biological) triplicate experiments. Results are expressed as mean ± SD (n = 3). ES-LONG: human ESC-derived mesenchymal stromal cells, ES-RYU: human ESC-derived intestinal epithelial cells, iPS-LONG: human iPSC-derived mesenchymal stromal cells, iPS-RYU: human iPSC-derived intestinal epithelial cells. (B) Induction of the cytochrome P450 genes with exposure to Omeprazole (OM), β-Naphthoflavone (BNF), Phenobarbital (PB), Rifampicin (Rif), vitamin D (VD3), and Dexamethasone (Dex). Results are expressed as mean ± SD (n = 3). The expression level of each gene without any treatment (DMSO) was set to 1.0. Each expression level was calculated from the results of independent (biological) triplicate experiments. Statistical significance was determined using Student’s t-test; ** Fold > 2, p < 0.01, * Fold > 2, p < 0.05. (C) Immunocytochemistry of the intestinal epithelial cells with an antibody to CYP3A4 and CES2, drug-metabolizing enzymes. Nuclei were stained with DAPI. Scale bars: 50 µm. (D) Measurement of CYP3A4 activity. The levels of 6β-hydroxy testosterone were measured. Results are expressed as mean ± SD (n = 3 triplicate biological experiments). Statistical significance was determined using Dunnett’s test compared to Caco-2.

To evaluate CYP3A4 activity, the intestinal epithelial cells were incubated with testosterone, and the formation of 6β-hydroxytestosterone, a metabolite produced from testosterone by CYP3A4, was measured. The CYP3A4 metabolic activity levels in the intestinal epithelial cells were 22-34-fold higher than those in Caco-2 cells (Figure 6D and S6E).

### Puromycin-based selection of intestinal epithelial cells

We then aimed to validate the impact of mesenchymal stromal cells by eradicating these cells through the use of a cytocidal antibiotic, puromycin.^19,23^ Puromycin is capable of preferentially selecting cells that possess drug-metabolizing enzymes while allowing for the survival of only intestinal epithelial cells (Figure S7A and B). After puromycin selection, the intestinal epithelial cells continued to exhibit positivity for ECAD, CDH17, ZO-1, Claudin-2 and Claudin-7, which are elements of adherens and tight junctions. We then evaluated the barrier function of the puromycin-treated intestinal epithelial cells by TEER measurements and Lucifer yellow assays (Figure S7D and E). The TEER value for the intestinal epithelial cells was 195-236 Ω cm² (Figure S7D). The Lucifer yellow Papp value was 0.67-1.41 x 10□□ cm/s (Figure S7E). These results demonstrate that the barrier function of intestinal epithelial cells is increased by placing intestinal epithelial cells on mesenchymal stromal feeders (Figure 5C, 5D, S7D and S7E). Subsequently, we followed up the time course of intestinal epithelial cells’ barrier function for 20 days (Figure S7F, S7G and S7H). Surprisingly, the puromycin-treated intestinal epithelial cells exhibited an unstable barrier function (Papp value: 2.3 - 6.0). We also performed a transport assay using Rhodamine123 in the puromycin-treated intestinal epithelial cells (Figure S7I). The puromycin-treated intestinal epithelial cells demonstrated asymmetric permeability, with efflux ratios (ER) 5.9-18.1. The puromycin-treated intestinal epithelial cells expressed transporters, P-gp, BCRP, PEPT1, OATP2B1, and MRP3, whereas the expression of MRP2 was limited (Figure 5H and S7J). These results suggest that mesenchymal stromal cells are required for long-term maintenance of intestinal epithelial cells.

## Discussion

This study shows that mesenchymal stromal cells derived from intestinal organoids play an important role in the proliferation and maintenance of intestinal epithelial cells in adherent cultures. This culture system is a valuable and readily available model for examining the niche environment such as the extracellular matrix and cytokines provided by mesenchymal cells. Organoid-derived mesenchymal stromal cells contribute to the early maturation and maintenance of the function of intestinal epithelial cells.

Intestinal organoids derived from LGR5+ stem cells require the presence of EGF, Noggin to counteract BMP signaling, as well as R-Spondin, a WNT agonist, and Wnt for maintenance.^7^ Mesenchymal stromal cells located in the intestinal mucosa exert a stimulatory effect on the intestinal epithelium.^12,41–43^ The contribution of mesenchymal stromal cells is a multifactorial process, not attributable to a single element, and mediated through the secretion of soluble factors and the provision of scaffolds. Epithelial cells are modulated by a concentration gradient of trophic factors, namely WNT and BMP signaling. WNT, R-Spondin, and BMP inhibitors are highly concentrated at the base of the crypts, where small intestinal stem cells reside, while the levels of these trophic factors decrease as the cells ascend to the villi.^41,44,45^ Ultimately, the predominance of the BMP ligand determines the fate of the cells. PDGFRA+ and CD81+ stromal cells (trophocytes) around the crypts produce a BMP inhibitor, GREM1. Mesenchymal stromal cells derived from intestinal organoids in the present study may contribute to intestinal epithelial cell proliferation through soluble factors such as BMP inhibitor GREM1.^18^ The mesenchymal stromal cells can be considered trophocytes, as they produce GREM1, an important component of the stem cell niche.

Intestinal epithelial stem cells are located at the base of the crypts. Intestinal stem cells exhibit specific markers such as LGR4+ and LGR5+, and LGR5 expression is lost upon differentiation.^29^ Enterocytes and goblet cells in the intestinal epithelial cell population probably suggest the presence of intestinal epithelial stem cells. High expression of LGR4 and a lack of LGR5 indicate that the intestinal epithelium with proliferative capacity and multipotency correspond to transient amplifying progenitor cells (TA cells). The CD24+/CD44+ intestinal epithelial cells in this study can be consistent with the LGR5-negative multipotent cell population at the base of the intestinal crypts.^27,28^ Intense Wnt signal-dependent multipotency and proliferation of TA cells may explain the strong proliferative potential of the organoid-derived intestinal epithelial cells (RYU) generated in this study.^27,46,47^

The intestinal epithelial cells on mesenchymal feeders possess barrier ability and can be applied in developing an absorption-accelerating agent. Caco-2 cells lack a mucosal layer,^48^ whereas pluripotent stem cell-derived intestinal epithelial cells have goblet cells and form a mucosal layer. Comparable barrier function and expression of P-gp and CYPs in the organoid-derived epithelial cells (RYU) with frozen human intestines suggest that the organoid-derived epithelial cells can contribute to drug development and pharmacokinetics. Furthermore, high expression of AHR, PXR, VDR, and GR and the induction of CYPs by the drugs through these receptors may lead to the availability of the cells as an in vitro simulator for drug-drug interactions.^36^ The inducibility of CYP3A by rifampicin has been documented in human intestinal enterocytes, whereas Caco-2 cells exhibit no such inducibility.^49^ The development of an accurate and reproducible differentiation method can contribute to the evaluation of intestinal absorption of drugs in humans. Because of the limited availability of cell types enabling the concurrent assessment of metabolic enzymes, transporters, and mucosal layers, intestinal epithelial cells can be a compelling model for evaluating the pharmacokinetics of the human small intestine. Moreover, the intestinal epithelial cells have the potential to reflect enterocyte functionality compared to Caco-2 cells model more accurately.

By our calculations based on the results, intestinal epithelial cells can proliferate at least up to 1 x 10^10^. Intestinal epithelial cells can form cell sheets and possibly contribute to the improvement of esophageal stricture as well as oral mucosal epithelial cells.^50^ Likewise, the cell sheets of intestinal epithelial cells improve inflammatory bowel diseases such as Crohn’s disease.^51^ Mesenchymal stromal cells in the sheets may mitigate inflammation through immune tolerance and anti-inflammatory cytokines like marrow stromal cells and adipocytes.^52^

## Conclusion

This study shows that mesenchymal stromal cells from self-assembled intestinal organoids generate functional enterocytes and LGR4-positive functional intestinal epithelial stem cells in two-dimensional culture. The combination of these epithelial stem cells and mesenchymal stromal cells achieves epithelial multipotency, precise lineage control by the stroma, a niche environment provided by the stroma, and self-organization of the epithelium. It also demonstrates the potential for building a robust platform for drug discovery and toxicology to study the effects of environmental pollutants, chemicals, and pharmaceuticals on humans. Furthermore, their proliferation rate, which is several orders of magnitude faster than organoids, has potential applications in regenerative medicine, preclinical studies, and disease models.

## MATERIAL AND METHODS

### Ethical statement

All experiments handling human cells and tissues were approved by the Institutional Review Board at the National Center for Child Health and Development (2021-178). Informed consent was obtained from all tissues and cell donors. When the donors were under 18, informed consent was obtained from parents. Human cells in this study were utilized in full compliance with the Ethical Guidelines for Medical and Health Research Involving Human Subjects (Ministry of Health, Labor, and Welfare (MHLW), Japan; Ministry of Education, Culture, Sports, Science and Technology (MEXT, Japan). The derivation and cultivation of human embryonic stem cell (hESC) lines were performed in full compliance with “the Guidelines for Derivation and Distribution of Human Embryonic Stem Cells (Notification of MEXT, No. 156 of August 21, 2009; Notification of MEXT, No. 86 of May 20, 2010) and “the Guidelines for Utilization of Human Embryonic Stem Cells (Notification of MEXT, No. 157 of August 21, 2009; Notification of MEXT, No. 87 of May 20, 2010)”. All procedures of animal experiments were approved by the Institutional Animal Care and Use Committee in National Center for Child Health and Development, based on the basic guidelines for the conduct of animal experiments in implementing agencies under the jurisdiction of the Ministry of Health, Labour and Welfare (Notification of MHLW, No. 0220-1 of February 20, 2015).

### Culture of human iPSCs

Human iPSCs (EDOM22 #8) derived from menstrual blood were used.^53,54^ The iPSCs were cultured on dishes coated with vitronectin (A14700, Thermo Fisher Scientific) in Stem flex medium (A3349401, Thermo Fisher Scientific). The dishes were coated with 5 μg/ml vitronectin at room temperature for 1 h. The medium was changed daily. The iPSCs were detached with 0.5 mM EDTA (06894-14, Nacalai tesque) for 5 min and passaged at 1,100 cells/cm^2^ every week.

### Culture of human ESCs

Human ESCs (SEES2) were cultured on dishes coated with iMatrix-511 silk (892021, nippi) in StemFit medium (RCAK02N, Ajinomoto).^55,56^ The dishes were coated with 1.7 μg/ml iMatrix-511 silk at 37°C for 1 h. The medium was changed daily. ESCs were detached with a 1:1 mixture of 0.5 mM EDTA and TrypLE™ Select (A1217701, Thermo Fisher Scientific) for 5 min and passaged at 550 cells/cm^2^ every week.

### Generation of intestinal organoid

To generate intestinal organoids, ESCs or iPSC were dissociated using 0.5 mM EDTA or a 1:1 mixture of 0.5 mM EDTA and TrypLE™ Select and plated on a cell-patterning glass substrate CytoGraph (Dai Nippon Printing Co. Ltd.). CytoGraphs were coated with 5 μg/ml vitronectin at room temperature for 30 min. The cells were cultured in XF32 medium (81% Knockout DMEM (10829018, Thermo Fisher Scientific), 15% Knockout Serum Replacement XF CTS (12618013, Thermo Fisher Scientific), 2 mM GlutaMAX (35050061, Thermo Fisher Scientific), 0.1 mM NEAA (11140050, Thermo Fisher Scientific), Penicillin-Streptomycin (15070063, Thermo Fisher Scientific), 50 μg/ml L-ascorbic acid 2-phosphate (A8960, Sigma-Aldrich), 10 ng/ml heregulin-1β (080-09001, Fujifilm Wako Pure Chemicals Co., Ltd.), 200 ng/ml recombinant human IGF-1 (85580C, Sigma-Aldrich), and 20 ng/ml human bFGF (PHG0021, Thermo Fisher Scientific)). The first day of the passage was performed with medium containing Rho-associated protein kinase inhibitor Y-27632 (036-24023, Fujifilm Wako Pure Chemicals Co., Ltd.). Gut-like peristaltic organoids were collected at 40 to 60 days after the passage and transferred to 6-well Ultralow adhesion plates (3471, Corning).

### Preparation of mouse embryonic fibroblasts

Mouse embryonic fibroblasts (MEF) were prepared for use as nutritional support (feeder) cells. Heads, limbs, tails, and internal organs were removed from E12.5 ICR mouse fetuses (Japan CLEA), and the remaining torsos were then minced with a blade and seeded into culture dishes with DMEM (D6429, Sigma-Aldrich) supplemented with 10% FBS (10091148, Thermo Fisher Scientific) to allow cell growth. After 2 days of culture, the cells were passaged in a 1:4 ratio. After 5 days of culture, cells were trypsinized with 0.25% trypsin/1 mM EDTA (209-16941, Fujifilm Wako Pure Chemicals Co., Ltd.) and 1/100 (v/v) of 1 M HEPES buffer (15630–106, Thermo Fisher Scientific) was added to the collected cells. Following irradiation with an X-ray apparatus (dose: 30 Gy, MBR-1520 R-3, Hitachi), the cells were cryopreserved with a TC protector (TCP-001DS, Pharma Biomedical, Osaka, Japan).

### Preparation of intestinal stromal feeder cells (LONG)

Intestinal organoids were cut and spread on a 35-mm dish coated with iMatrix-511 silk. The dishes were coated with 1.7 μg/ml iMatrix-511 silk at 37°C for 1 h. Then the attached fragments were cultured at 37°C in 5% CO2 with ESTEM-HE medium (GlycoTechnica). The medium was changed every 2 days. After 21 days of culture, the cells were trypsinized with 0.25% trypsin/1 mM EDTA for 3 min. The passage was onto a non-coated plate. The cells were cultured in XF32 medium at 0-3 days and then in ESTEM-HE medium at day 4 until just before passage. This passaging was performed at least three times, and the cells were purified before use for feeder cells. Thereafter, the medium was changed to DMEM supplemented with 10% FBS at 37°C in 5% CO2. The medium was changed every 3 days. The cells were passaged every 5-7 days and seeded at 3.6 x 10^4^ cells/cm^2^. The cells were cryopreserved with STEM-CELLBANKER (ZR646, Nippon Zenyaku Kogyo Co., Ltd.) until use. The protocol for intestinal mesenchymal stromal cell production is shown in a flow chart (Figure S4A).

### Preparation of intestinal epithelial cells (RYU)

Intestinal organoids were cut and spread on a 35-mm dish coated with iMatrix-511 silk at 37°C for 1 h. Dishes were coated with 1.7 μg/ml iMatrix-511 silk at 37°C for 1 h. Then, fragments that had attached were cultured at 37°C in 5% CO2 with ESTEM-HE medium. The medium was changed every 2 days. After 28 days of culture, the cells were detached with 0.25% trypsin/1 mM EDTA and seeded onto confluent intestinal mesenchymal stromal cells. The cells were then cultured at 37°C in 5% CO2 in ESTEM-HE medium. The medium was changed every 2 days. The cells were passaged into 4-8 dishes after detachment with 0.25% trypsin/1 mM EDTA. The cells were cryopreserved with STEM-CELLBANKER. The procedure is shown in flow charts (Figure S4A).

### Preparation for histological studies

Samples were coagulated in iPGell (PG20-1, GenoStaff) following the manufacturer’s instructions and fixed in 4% paraformaldehyde at 4°C overnight. Fixed samples were embedded in a paraffin block to prepare thin cell sections. Deparaffinization, dehydration, and permeabilization were performed using standard techniques. Hematoxylin-eosin (HE) staining was performed with Carrazzi’s hematoxylin solution (30022, Muto Chemicals) and eosin Y (32053, Muto Chemicals). Alcian blue staining was achieved with alcian blue solution pH2.5 (40852, Muto Chemicals). Nuclei were counterstained with kernechtrot solution (40872, Muto chemicals).

### Immunostaining

Cells in culture (3910-035, IWAKI) were fixed with 4% paraformaldehyde for 15 min at room temperature. After washing with phosphate-buffered saline (PBS), cells were permeabilized with 0.25% Triton X-100 in PBS for 20 min, pre-incubated with Protein Block Serum-Free (X0909, Dako) for 30 min at room temperature and then exposed to primary antibodies overnight at 4 °C. After three washes with PBS, the cells were incubated with fluorescent secondary antibodies for 1 h at room temperature and washed three times with PBS. Nuclei were stained with a mounting medium containing 4’,6-diamidino-2-phenylindole dihydrochloride solution (DAPI) (H-1200, Vector Laboratories).

For immunofluorescence staining of thin cell sections, deparaffinized cell sections were unmasked epitopes by heating with histophine (415211, NICHIREI) in a microwave oven at a power setting of 700 W for 20 min. After three washes with PBS, sections were incubated with primary antibodies overnight at 4°C. After three washes with PBS, the sections were incubated with secondary antibodies at room temperature for 30 min and washed three times with PBS. Nuclei were counterstained with a mounting medium containing DAPI (H-1800, Vector Laboratories).

When peroxidase-conjugated secondary antibodies were employed (424154, NICHIREI), cell sections were exposed to 3% hydrogen peroxide/methanol for 5 min to block endogenous peroxidase and color development was performed with 3,3’-diaminobenzidine (4065-1, Muto Chemicals). The sections were counterstained with hematoxylin.

The antibodies listed in Tables 1 and 2 were diluted as indicated in PBS containing 1% BSA (126575, Calbiochem).

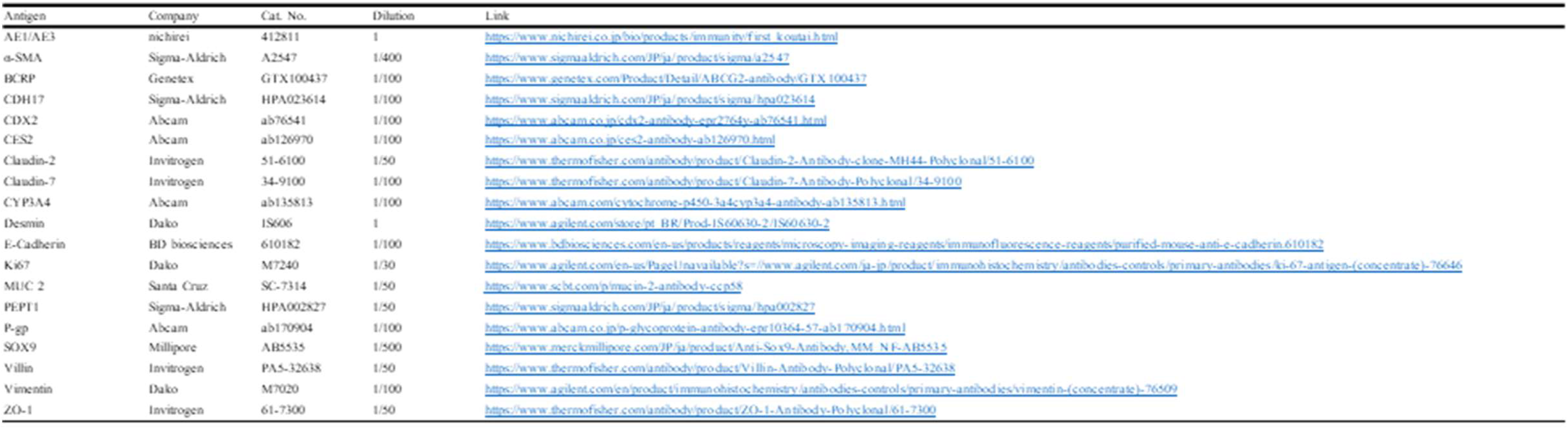
Table 1.

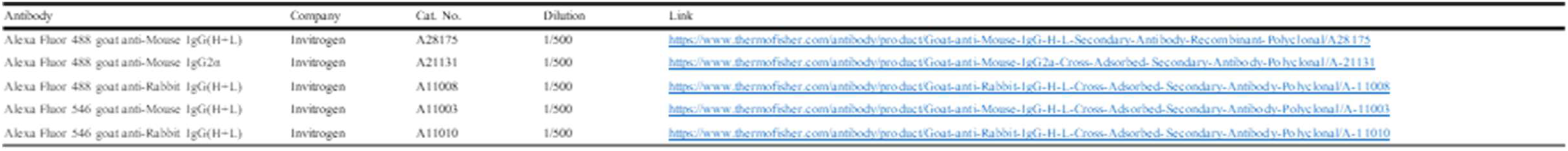
Table 2.

Table 2
| Antibody | Company | Cat. No. | Dilution | Link |
| --- | --- | --- | --- | --- |
| Alexa Fluor 488 goat anti-Mouse IgG(H+L) | Invitrogen | A28175 | 1/500 | https://www.thermofisher.com/antibody/product/Goat-anti-Mouse-IgG-H-L-Secondary-Antibody-Recombinant-Polyclonal/A28175 |
| Alexa Fluor 488 goat anti-Mouse IgG2a | Invitrogen | A21131 | 1/500 | https://www.thermofisher.com/antibody/product/Goat-anti-Mouse-IgG2a-Cross-Adsorbed-Secondary-Antibody-Polyclonal/A-21131 |
| Alexa Fluor 488 goat anti-Rabbit IgG(H+L) | Invitrogen | A11008 | 1/500 | https://www.thermofisher.com/antibody/product/Goat-anti-Rabbit-IgG-H-L-Cross-Adsorbed-Secondary-Antibody-Polyclonal/A-11008 |
| Alexa Fluor 546 goat anti-Mouse IgG(H+L) | Invitrogen | A11003 | 1/500 | https://www.thermofisher.com/antibody/product/Goat-anti-Mouse-IgG-H-L-Cross-Adsorbed-Secondary-Antibody-Polyclonal/A-11003 |
| Alexa Fluor 546 goat anti-Rabbit IgG(H+L) | Invitrogen | A11010 | 1/500 | https://www.thermofisher.com/antibody/product/Goat-anti-Rabbit-IgG-H-L-Cross-Adsorbed-Secondary-Antibody-Polyclonal/A-11010 |

### Population Doubling

Cells were seeded from one 10-cm dish (353003, Falcon) into four 10-cm dishes at each passage. Population doubling was estimated from passage number.

### Karyotypic analysis

Karyotypic analysis was contracted out to Nihon Gene Research Laboratories (Sendai, Japan). To assess diploidy, 50 cells at metaphase were examined. Metaphase spreads were prepared from cells treated with 100 ng/mL of Colcemid (KaryoMax, Gibco; Thermo Fisher Scientific, MA, USA) for 6 h. The cells were fixed with methanol: glacial acetic acid (2:5) three times and placed onto glass slides. Giemsa banding was applied to metaphase chromosomes. A minimum of 10 metaphase spreads were analyzed for each sample and karyotyped using a chromosome imaging analyzer system (Applied Spectral Imaging, CA, USA).

### Microarray analysis

Total RNA was isolated using miRNeasy mini kit (217004, Qiagen, Hilden, Germany). RNA samples were labeled and hybridized to a SurePrint G3 Human GEO microarray 8□×□60 K Ver 3.0 (Agilent, CA, USA), and the raw data were normalized using the 75-percentile shift. Gene expression profiles of hSI were analyzed using human small intestine total RNA (R1234226-50, biochain). Results were visualized with cluster maps and boxplots using the R package ggplot2 and ComplexHeatmap.

### Quantitative RT-PCR analysis

The cultured cells underwent two washes with Dulbecco’s phosphate-buffered saline (Sigma-Aldrich, St Louis, MO, USA) before the isolation of total RNA. RNA extraction was carried out using the RNeasy total RNA extraction kit (Qiagen, Hilden, Germany) as per the manufacturer’s guidelines. To quantify the gene expression levels, qPCR was performed using 8 ng of total RNA, following reverse transcription with the High Capacity RNA-to-cDNA kit (Thermo Fisher Scientific, Waltham, MA, USA) according to the manufacturer’s protocol. The QuantStudio 7 Flex Real-Time PCR system (Applied Biosystems, Foster City, CA, USA) was used to measure gene expression levels, and primers and probe sets were utilized to detect each gene transcript, as listed in Table 3. The expression levels of the genes are presented relative to RNAs derived from a human small intestine (R1234226-50, BioChain Institute, Inc., Newark, CA, USA).

Table 3.

Table 3
| Gene Symbole | Assay ID |
| --- | --- |
| CYP1A1 | Hs01054797_g1 |
| CYP1A2 | Hs00167927_m1 |
| CYP2B6 | Hs04183483_g1 |
| CYP2C9 | Hs00426397_m1 |
| CYP2C19 | Hs00426380_m1 |
| CYP2D6 | Hs00164385_m1 |
| CYP3A4 | Hs00430021_m1 |
| VDR | Hs00172113_m1 |
| PXR | Hs00243666_m1 |
| AHR | Hs00169233_m1 |
| GR | Hs00230813_m1 |
| P-gp | Hs00184500_m1 |
| BCRP | Hs01053790_m1 |
| PEPT1 | Hs00192639_m1 |
| OATP2B1 | Hs01030343_m1 |
| MRP2 | HS00960489_m1 |
| MRP3 | Hs00978452_m1 |

### Cell transplantation

Male immunodeficient (NOD.Cg-PrkdcscidIl2rgtm1Sug/ShiJic) mice, aged 6–8 weeks, (Charles River Laboratories, Inc., Wilmington, MA) were used in this study. The mice were anesthetized by inhalation of 3% isoflurane (099-06571, Fujifilm Wako Pure Chemicals Co., Ltd.). Under sterile conditions, a 1–2□cm incision was made below the ribs and 0.5□cm off-center but parallel to the spine. One kidney was gently pulled out and kept moist with sterile saline. A cell suspension was gently pulled into a 1 ml syringe with a 27G needle. The injection site was located at the upper or/and lateral side of the kidney to avoid damage to kidney blood vessels. The needle was slowly pushed into the capsule, as far as possible, toward the inferior pole of the kidney to avoid perforating the kidney capsule in other areas. The cells in 50□μL of medium were injected into the kidney capsule with a syringe, a sterile cotton swab was placed over the kidney capsule to prevent cell leakage, and the needle was slowly withdrawn. The kidney was put back into the body cavity using the sterile cotton swab without applying pressure to it. Finally, the abdominal wall was closed with a suture.

### Transmission electron microscopy (TEM) and scanning electron microscopy (SEM)

For transmission electron microscopy of cultured cells, cells were washed three times with PBS. Fixation was performed in PBS containing 2.5% glutaraldehyde for 2 h. The cells were embedded in epoxy resin. Ultrathin sections cut vertically to the culture surface were double stained in uranyl acetate and lead citrate and were viewed under a JEM-1200 PLUS transmission electron microscope (Nihon Denshi) and SU6600 (Hitachi High-Tech corporation).

### Caco-2 Cell Culture

Caco-2 cells were cultured DMEM supplemented with 10% FBS at 37°C in 5% CO2 for 14 days. The medium was changed every 2 days. The cells were seeded at 2.0 x 10^5^ cells/cm^2^.

### Transepithelial electrical resistance (TEER) measurements

The cells were seeded onto Transwell inserts (353095, Falcon) and cultured until confluent. The Millicell-ERS (Electrical Resistance System) with chopstick electrodes was used to measure construct transepithelial electrical resistance (TEER). The instrument was calibrated with a 1000 Ω resistor prior to measurements, and an empty Transwell was used as a blank. Constructs were equilibrated to room temperature and switched to basal media for readings. The blank Transwell and all samples were measured three times. Samples and blanks were measured in triplicate, averaged, and used in the following calculations:

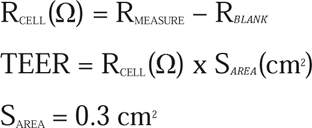

### Permeability and transport assay

The cells were seeded on the inserts of Transwells and cultured until confluent. Before the start of the assay, samples were pre-incubated with transport buffer (HBSS (14025092, Gibco) supplemented with 10 nM HEPES (15630106, Gibco) and 4.5 mg/ml glucose) for 1 h. Lucifer yellow solution (125-06281, Fujifilm Wako Pure Chemicals Co., Ltd.) was added to the apical side to a final concentration of 300 μM and then drawn out of the basal side at 30 min intervals for up to 120 min at 37°C with shaking at 40 rpm. Rhodamine123 solution (187-01703, Fujifilm Wako Pure Chemicals Co., Ltd.) was added to the apical or basal side to a final concentration of 5 μM and then drawn out of the basal side at 60 min intervals for up to 120 min at 37°C with shaking 40 rpm. An equal volume of transport buffer was immediately added to the drawn out side after each sampling. Lucifer yellow fluorescence was measured using a SYNERGY H1 microplate reader (Bio-Tek) at 428 nm excitation and 536 nm emission, and Rhodamine123 fluorescence was measured at 480 nm excitation and 530 nm emission. Papp was calculated using the following equation:

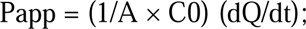

where dQ/dt is the rate of drug permeation across the cell monolayer, C0 is the donor-compartment concentration at time zero and A is the area of the cell monolayer. The ER was defined as Papp (basal-to-apical)/Papp (apical-to-basal).

### Cytochrome P450 induction assays

Cells cultured in the 24-well plates in ESTEM-HE medium were treated with 100 nM vitaminD3 (D1530-10UG, Sigma-Aldrich)or 40 μM β-naphthoflavone (N3633-1G, Sigma-Aldrich) or 50 μM omeprazole (150-02091, Fujifilm Wako Pure Chemicals Co., Ltd.) for 24 h or 20 μM rifampicin (189-01001, Fujifilm Wako Pure Chemicals Co., Ltd.) or 500 μM phenobarbital(162-11602, Fujifilm Wako Pure Chemicals Co., Ltd.) or 100 μM dexamethasone (194561, MP Biomedicals, Illkirch, France) for 48 h at a cell density of 80%. Controls were treated with DMSO (final concentration 0.1%).

### Determination of CYP3A4 activity

The cells were incubated with the assay buffer (HBSS supplemented with 10 nM HEPES and 4.5 mg/ml glucose) containing 50 µM testosterone (T1500, Sigma-Aldrich) at 37°C for 2 h and the assay medium was withdrawn at 30, 60, 120 min. The cells were finally detached with 0.25% trypsin/1 mM EDTA and collected for cell counting. The amount of the testosterone metabolite, 6β′-hydroxylated, was measured by LC-MS/MS (InertSustain^®^ AQ-C18, LC-20A (SHIMADZU), API4000 (AB Sciex Pte. Ltd.)). The measurements were performed by Sumika Chemical Analysis Service, Ltd.

## Declarations

### Ethics approval and consent to participate

Human cells in this study were performed in full compliance with the Ethical Guidelines for Clinical Studies (Ministry of Health, Labor, and Welfare, Japan). The cells were banked after approval of the Institutional Review Board at the National Institute of Biomedical Innovation (May 9, 2006). Animal experiments were performed according to protocols approved by the Institutional Animal Care and Use Committee of the National Research Institute for Child Health and Development.

### Availability of data and material

The datasets and cells used during the current study are available from the corresponding author on reasonable request.

### Authors’ contributions

AU and CJ designed the experiments. CJ, SH, SK, HK, SA and TA performed the experiments. CJ, TKaw, SH and SA analyzed data. SH, SA, TKawa, TKiy, HA, and SI contributed to the reagents, tissues and analysis tools. CJ, SA, SH, TKim, TKiy, SI and AU discussed the data and manuscript. CJ and AU wrote this manuscript. All authors read and approved the final manuscript.

## Acknowledgments

We would like to express our sincere thanks to K. Miyado and M. Komura for fruitful discussion, to M. Ichinose for providing expert technical assistance, to C. Ketcham for English editing and proofreading, and to E. Suzuki and K. Saito for secretarial work.

## Declaration of generative AI and AI-assisted technologies in the writing process

During the preparation of this work, CJ and AU used DeepL, Grammarly, and ChatGPT for English writing. After using DeepL, Grammarly, and ChatGPT, CJ and AU reviewed and edited the content as needed and take full responsibility for the content of the publication.

## Supplementary Figure Legends

**Figure S1. Mesenchymal stromal cells supported intestinal epithelial cells**

(A) Histology and immunohistochemistry of intestinal epithelial cells on MEFs. Immunohistochemistry was performed with antibodies to CDX2, Villin, and AE1/AE3. Scale bars: 200 µm (upper panels), 50 µm (lower panels). (B) Phase-contrast photomicrographs of mesenchymal stromal cells at Passages 5, 10, and 15. The mesenchymal stromal cells ceased their proliferation at Passage 15. Scale bars: 500 µm. (C) Histology and immunohistochemistry of intestinal epithelial cells on mesenchymal stromal cells. Scale bars: 200 µm (upper panels), 50 µm (lower panels).

**Figure S2. Intestinal epithelial cells require mesenchymal stromal cells**

(A) Phase-contrast photomicrograph (left-upper) and histology (the others) of intestinal epithelial cells on mesenchymal stromal cells (LONG). (Right-upper) H.E. stain. (Lower) Immunohistochemistry of intestinal epithelial cells with antibodies to CDX2 (left-lower) and Villin (right-lower). Scale bars: 500 µm (left-upper) and 50 µm (the others). (B) Phase-contrast photomicrograph (left-upper) and histology (the others) of intestinal epithelial cells on the non-coated dish. (Right-upper) H.E. stain. (Lower) Immunohistochemistry of intestinal epithelial cells with antibodies to CDX2 (left-lower) and Villin (right-lower). Scale bars: 500 µm (left-upper) and 50 µm (the others). (C) Phase-contrast photomicrograph (left-upper) and histology (the others) of intestinal epithelial cells on the extracellular matrix-coated dish (Matrigel, Collagen type-I, IV, Laminin-511 and 521). (Right-upper) H.E. stain. (Lower) Immunohistochemistry of intestinal epithelial cells with antibodies to CDX2 (left-lower) and Villin (right-lower). Scale bars: 500 µm (left-upper) and 50 µm (the others).

**Figure S3. Mesenchymal stromal cells promote proliferation of the intestinal epithelial cell organoids**

(A) Phase-contrast photomicrographs of intestinal epithelial cell organoids on mesenchymal stromal cells at days 6, 8, and 11 in Matrigel. Left: Intestinal epithelial cell organoids on mesenchymal stromal cells. Right: Intestinal epithelial cell organoids alone (no feeder cells). Scale bars: 500 µm. (B) Phase-contrast photomicrographs of co-culture of intestinal epithelial cell organoids and mesenchymal stromal cells on mesenchymal stromal cells at day 6, 8, and 11 in Matrigel. Left: co-culture on mesenchymal stromal cells. Right: co-culture alone (no feeder cells). Scale bars: 500 µm.

**Figure S4. Generation of intestinal epithelial and mesenchymal stromal cells from intestinal organoids**

(A) Experimental scheme for generation of the intestinal epithelial and mesenchymal stromal cells from small intestinal organoids. Intestinal epithelial cells and mesenchymal stromal cells were isolated at 21 and 28 days, respectively. (B) Details of intestinal epithelial and mesenchymal stromal cells established from human iPSC- and ESC-derived intestinal organoids.

**Figure S5. Intestinal epithelial cells are positive for crypt base cell (CBC) markers such as LGR4, CD24, and CD44**

(A) Immunohistochemistry of intestinal epithelial cells in iPGell (upper panel) and human small intestine tissue (lower panel) with antibodies to SOX9. Scale bars: 50 µm. (B) Expression heat map of gene expression in the intestinal epithelial cells and mesenchymal stromal cells. n = 3 triplicate biological experiments. (C) Boxplot of representative human intestinal epithelial stem cells and differentiated cell markers. n = 3 triplicate biological experiments (intestinal epithelial cells and mesenchymal stromal cells). Boxplots are expressed as mean ± SD. (D) Immunohistochemistry of organoid-derived intestinal epithelial cells and mesenchymal stromal cells at the subrenal capsule of the kidney 7 days after the implantation, using antibodies to CDX2, AE1/3, and Vimentin. Scale bar of zoomed sections represents 100 µm.

**Figure S6. Pharmacokinetics-related gene expression in intestinal epithelial cells**

(A) Heat map of gene expression in intestinal epithelial cells (Passages 5 and 12) and mesenchymal stromal cells. Each expression level was calculated from the results of triplicate biological experiments. (B) Boxplot of representative human intestinal epithelial stem cells and differentiated cell markers. Boxplots are expressed as mean ± SD (n = 3 triplicate biological experiments). (C) Expression of the genes for nuclear receptors (VDR, PXR, AHR, and GR). The expression level of adult small intestine whole tissue was set to 1.0. Each expression level was calculated from the results of triplicate biological experiments. Results are expressed as mean ± SD (n = 3 triplicate biological experiments). ES-LONG: human ESC-derived mesenchymal stromal cells, ES-RYU: human ESC-derived intestinal epithelial cells, iPS-LONG: human iPSC-derived mesenchymal stromal cells, iPS-RYU: human iPSC-derived intestinal epithelial cells. (D) Inhibition and induction of the cytochrome P450 genes with exposure to Rifampicin (Rif) and Ketoconazole (Ket). Results are expressed as mean ± SD (n = 3). The expression level of each gene without any treatment (DMSO) was set to 1.0. Each expression level was calculated from the results of independent (biological) triplicate experiments. (E) 6β-hydroxy testosterone concentration over time in CYP3A4 activity tests. ES-RYU: human ESC-derived intestinal epithelial cells (blue), iPS-RYU: human iPSC-derived intestinal epithelial cells (orange), Caco-2 (gray).

**Figure S7. Puromycin-based selection of intestinal epithelial cells**

(A) Phase-contrast photomicrographs of the intestinal epithelial cells, mesenchymal stromal cells and human ESCs with exposure to puromycin. Puromycin (1 µg/mL) was added (Day 0). Scale bars: 500 µm. (B) Experimental schematic for intestinal epithelial cells on mesenchymal stromal cells. (C) Phase-contrast micrograph and immunocytochemistry of epithelial cells with exposure to puromycin. Immunocytochemistry was performed with the antibody to barrier functions (ECAD, CDH17, ZO-1, Claudin-2, Claudin-7) and CDX2. Nuclei were stained with DAPI. Scale bars: 500 (black) and 50 (yellow) µm. (D) Trans-epithelial electrical resistance measurements for the intestinal epithelial cells. Mesenchymal stromal cells and Caco-2 cells served as negative and positive controls, respectively. White bars are puromycin-untreated intestinal epithelial cells on mesenchymal stromal cells, and black bars are puromycin-treated intestinal epithelial cells on mesenchymal stromal cells. Results are expressed as mean ± SD (n = 3 triplicate biological experiments). Statistical significance was determined using Student’s t-test. (E) Lucifer Yellow permeability tests for the intestinal epithelial cells on mesenchymal stromal cells. Caco-2 cells served as a control. Black and white bars are treated and non-treated with puromycin, respectively. Results are expressed as mean ± SD (n = 3 triplicate biological experiments). Statistical significance was determined using Student’s t-test. *<0.05, **<0.01. (F) Lucifer Yellow permeability test was performed for barrier performance. Barrier performance of epithelial cells (Puromycin-treated (+) and non-treated (-)) was altered in culture. Blue line, Puromycin-treated; orange line, Puromycin-untreated. Results are expressed as mean ± SD (n = 3 triplicate biological experiments). Statistical significance was determined using Student’s t-test; *<0.05, **<0.01. (G) Lucifer yellow concentration over time in lucifer yellow permeability tests 16 and 20 days after confluence. Puromycin-untreated (-, orange), Puromycin-treated (+, blue). (H) Phase-contrast photomicrographs of intestinal epithelial cells on mesenchymal stromal cells (Puromycin-treated (+) and non-treated (-)) 16 and 20 days after confluence. Pre-and post-Lucifer Yellow permeability test is shown on the left and right panels, respectively. Scale bars: 500 µm. (I) Permeability test with Rhodamine 123. The efflux ratio (ER) for Rhodamine 123 was derived from the Papp values associated with basal-to-apical transport (B to A) and apical-to-basal transport (A to B). Intestinal epithelial cells derived from human iPSCs and ESCs on mesenchymal stromal cells. Analysis was performed in the Puromycin-treated (+) or non-treated (-). ESC-RYU: ESC-derived intestinal epithelial cells, iPSC-RYU: iPSC-derived intestinal epithelial cells. Caco-2 cells served a positive control. Results are expressed as mean ± SD (n = 3 triplicate biological experiments). (J) Expression of the genes for transporters (P-gp, BCRP, PEPT1, OATP2B1, MRP2, and MRP3). The expression level of adult small intestine whole tissue was set to 1.0. Each expression level was calculated from the results of independent (biological) triplicate experiments. Black bars are puromycin-untreated, and gray bars are puromycin-treated intestinal epithelial cells on mesenchymal stromal cells. Results are expressed as mean ± SD (n = 3). ES-RYU: human ESC-derived intestinal epithelial cells, iPS-RYU: human iPSC-derived intestinal epithelial cells.

## Supplemental materials

### Supplemental Video

Time-lapse image of the proliferation of ESC-derived intestinal epithelial cells on the mesenchymal stromal cells

