## Supplementary figures and images for "Human intestinal organoid-derived PDGFRα+ mesenchymal stroma empowers LGR4+ epithelial stem cells"

### Figure S1

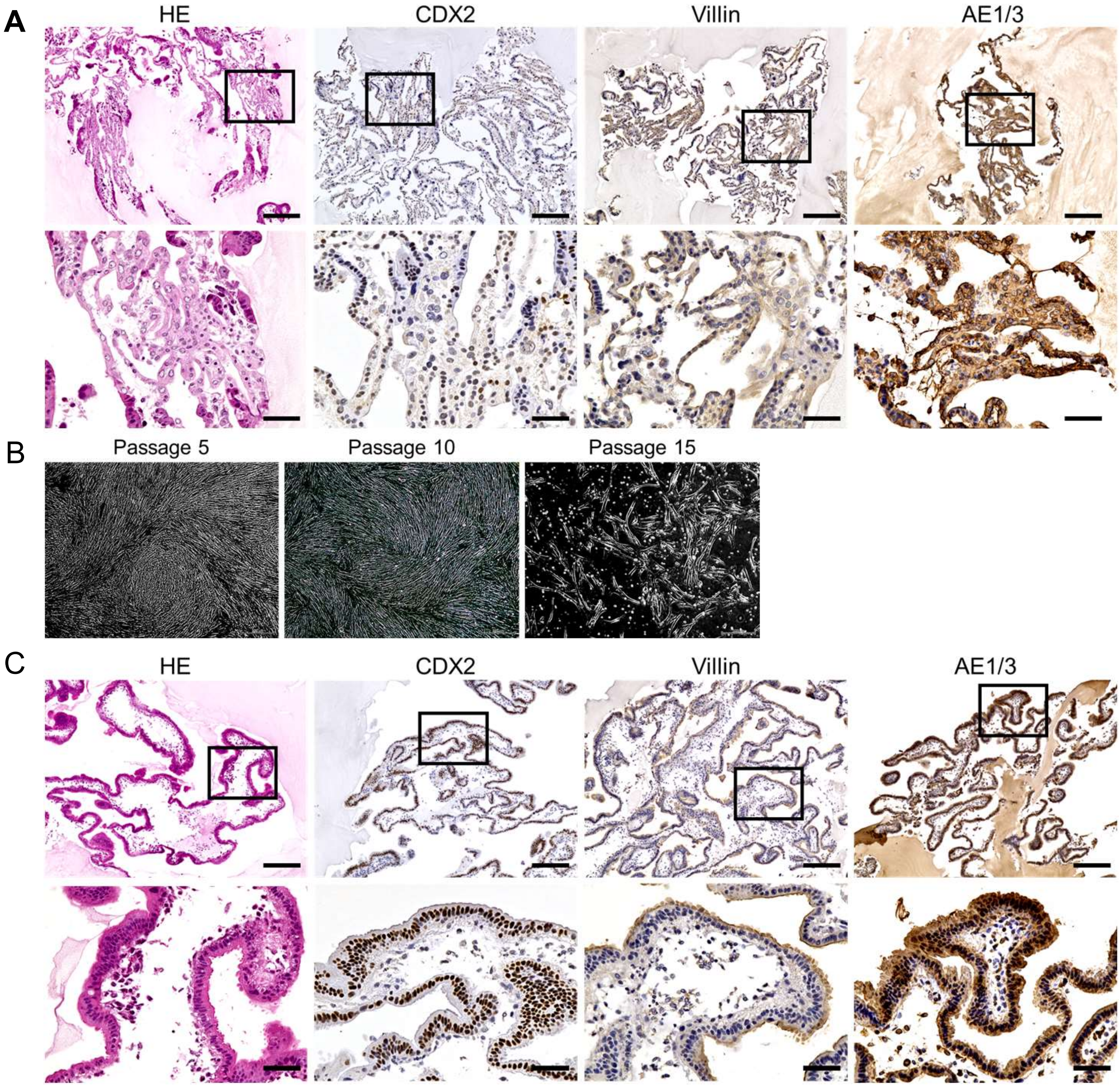


**Figure S1. Mesenchymal stromal cells generated intestinal epithelial cells**

### Figure S2

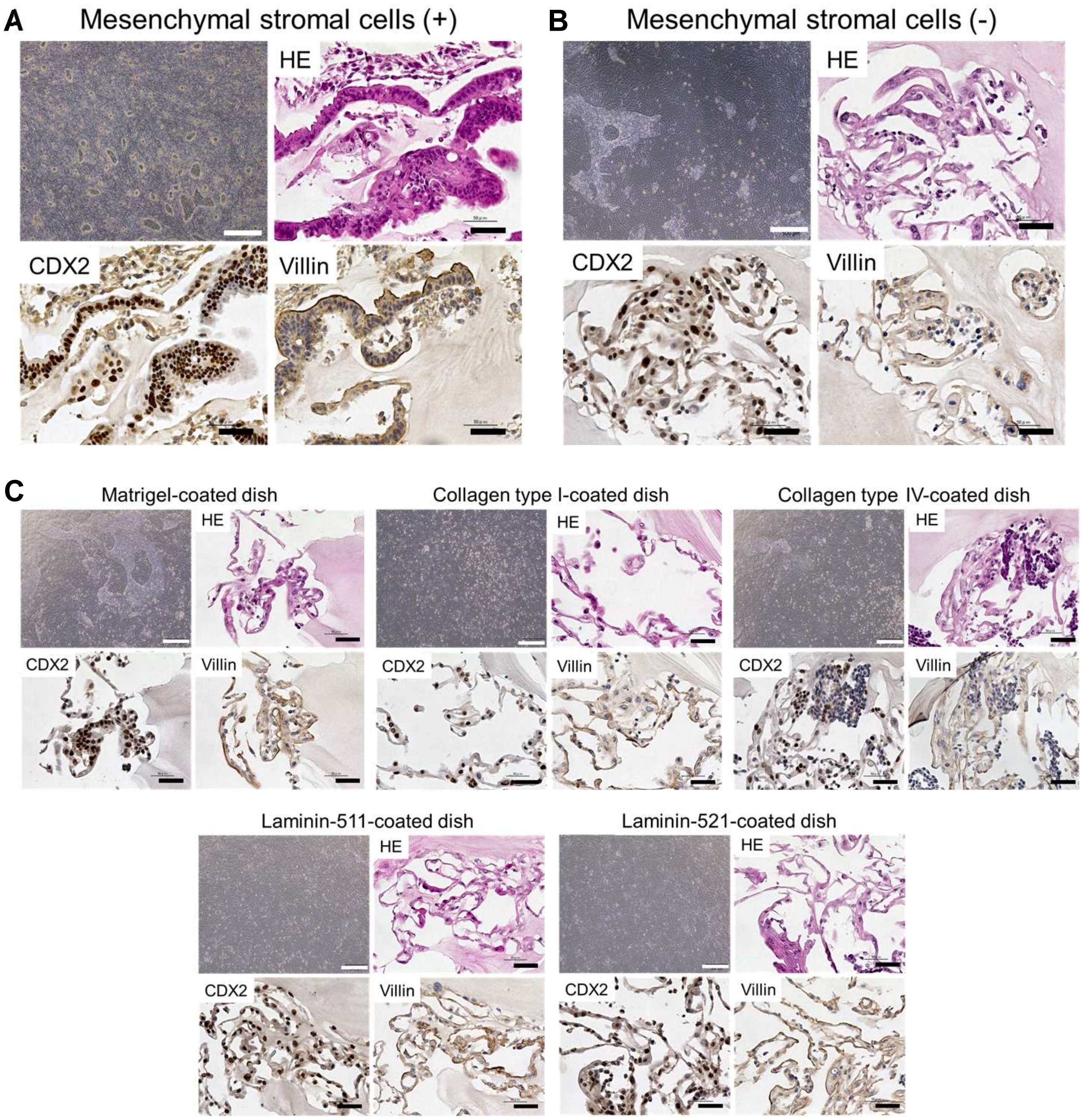


**Figure S2. Intestinal epithelial cells require mesenchymal stromal cells**

### Figure S6

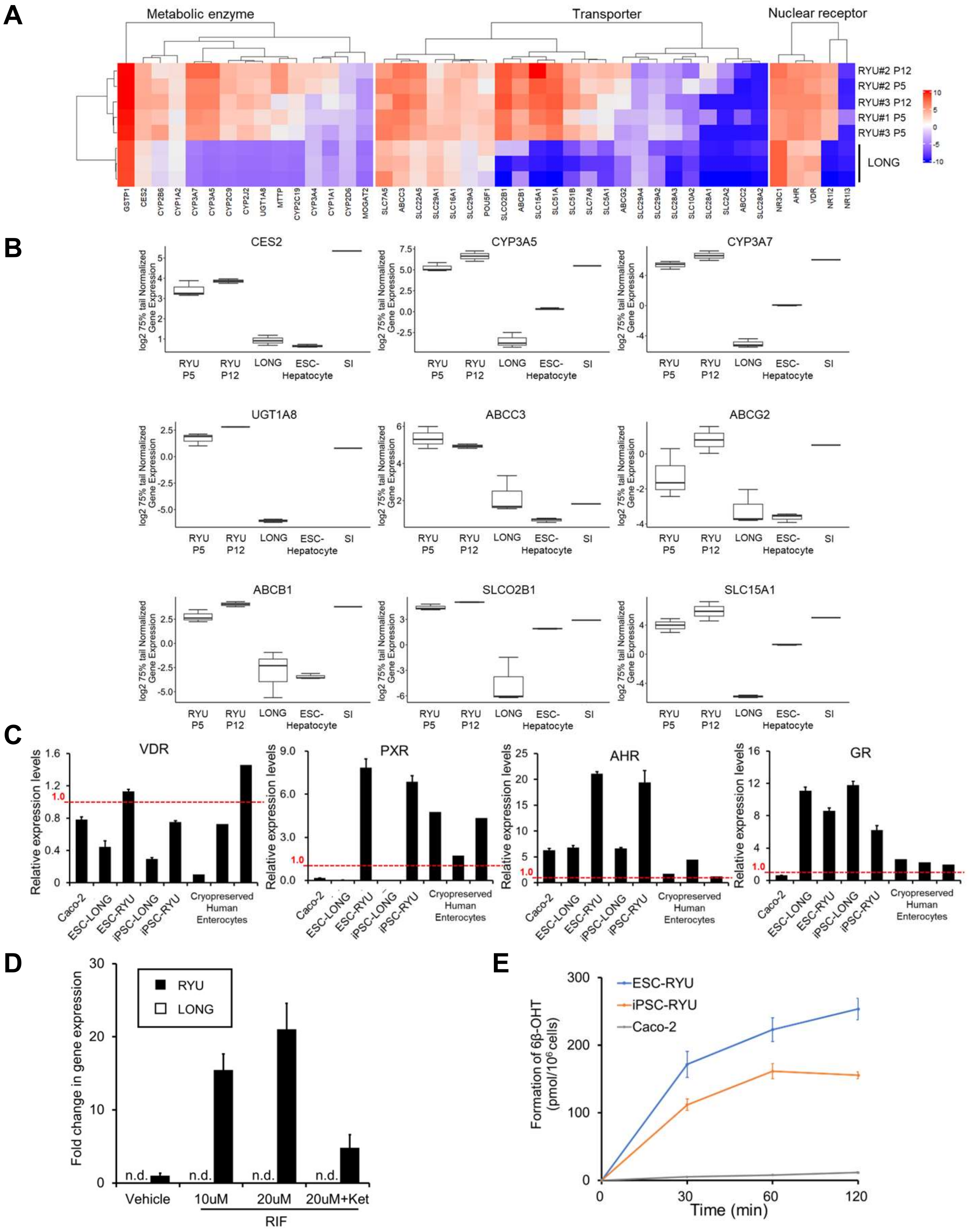


**Figure S6. Pharmacokinetics-related gene expression in intestinal epithelial cells**

### Figure S7

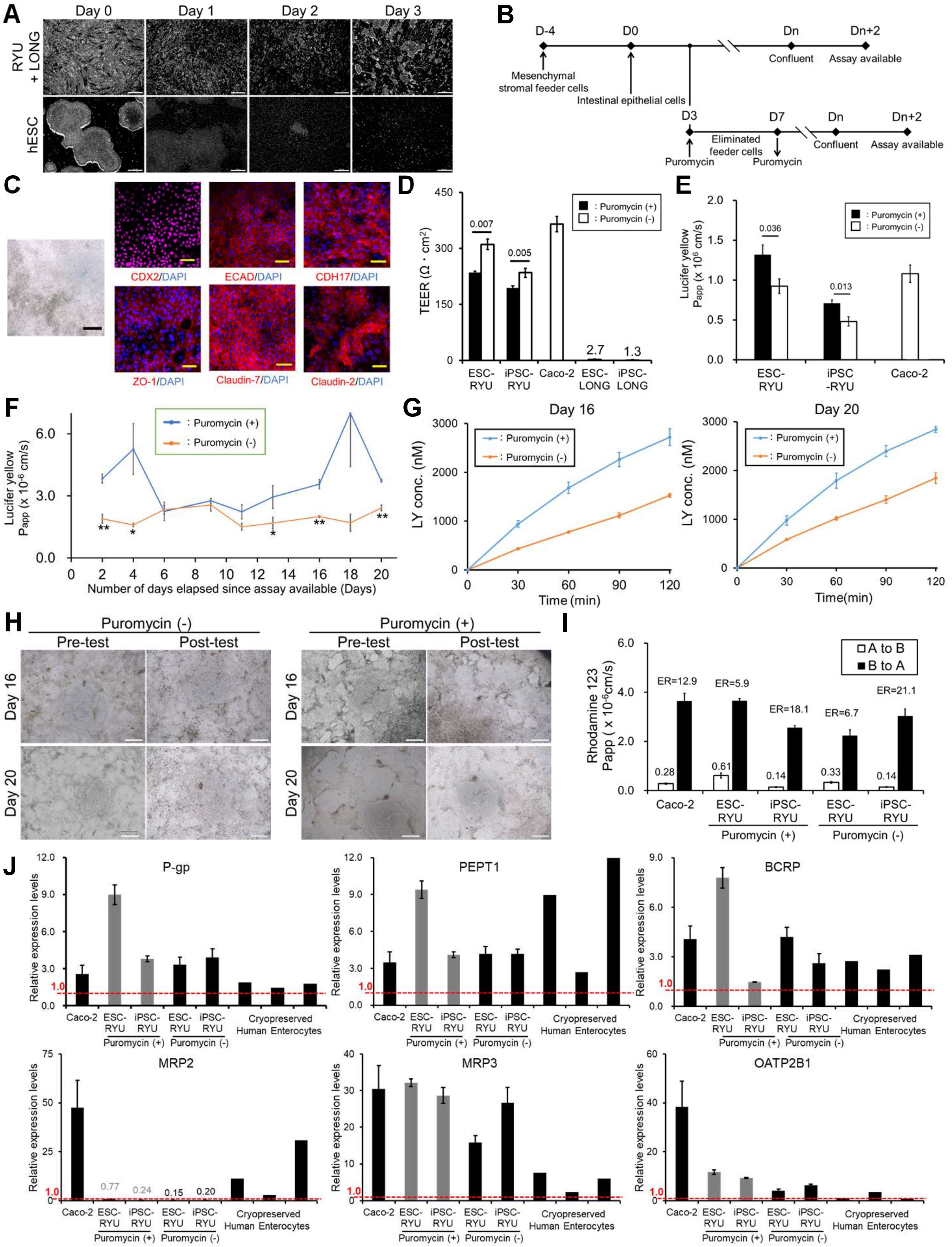


**Figure S7. Puromycin-based selection of intestinal epithelial cells**
