## Supplementary material for "Human intestinal organoid-derived PDGFRα+ mesenchymal stroma empowers LGR4+ epithelial stem cells": Figure S3

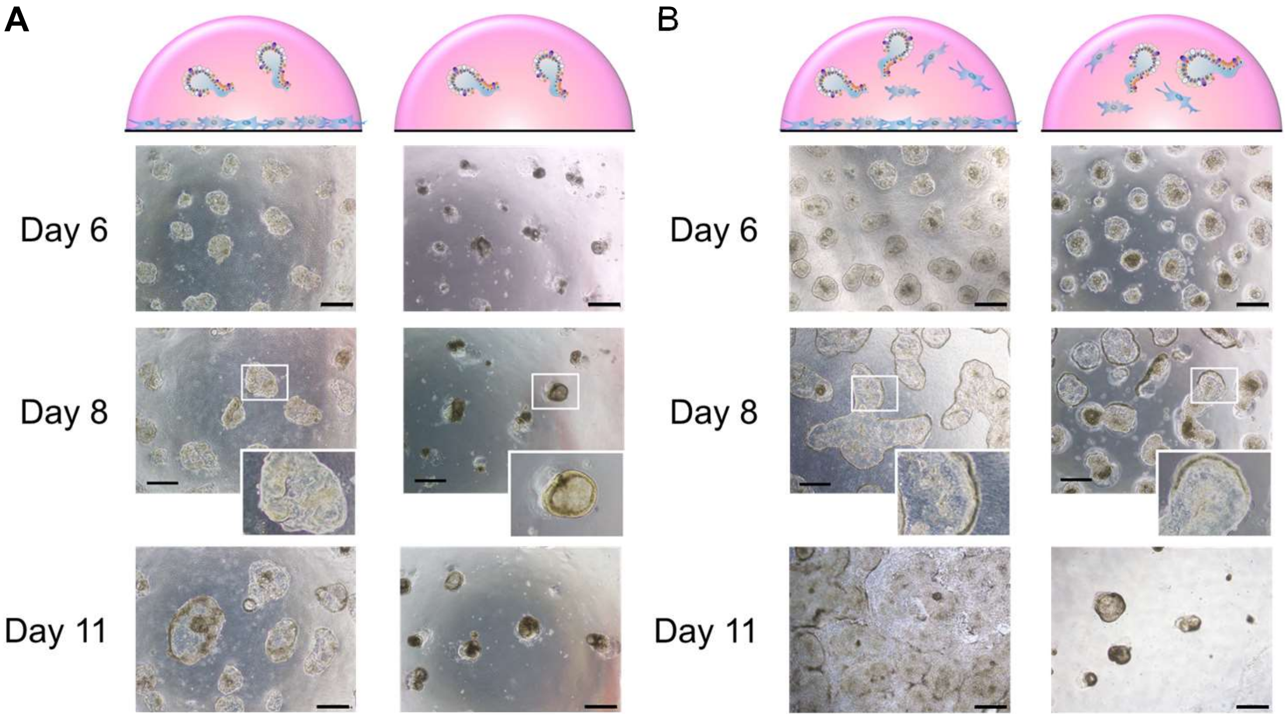


**Figure S3. Mesenchymal stromal cells promote proliferation of the intestinal epithelial cell organoids**
