## Supplementary material for "Human intestinal organoid-derived PDGFRα+ mesenchymal stroma empowers LGR4+ epithelial stem cells": Figure S4

A

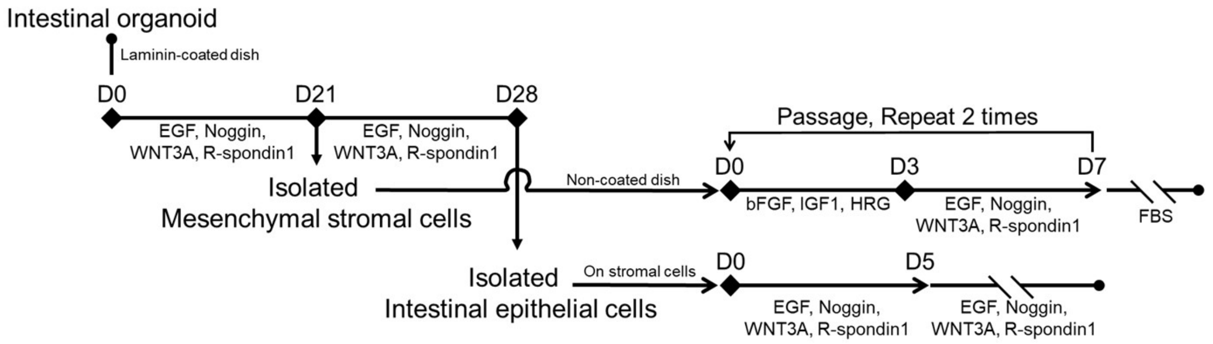


B

| Organoid Number | Origin | RYU Number | Established | LONG Number | Established | Condition |
|---|---|---|---|---|---|---|
| Intestinal organoid #1 | Human iPSCs | RYU01 | Success | LONG01 | Success | RYU：Normal , LONG：Normal |
| Intestinal organoid #2 | Human ESCs | RYU02 | Success | LONG02 | Success | RYU：Normal , LONG：Normal |
| | | RYU03 | Success | LONG03 | Success | RYU：Normal , LONG：Normal |
| Intestinal organoid #3 | Human ESCs | RYU04 | Success | LONG04 | Success | RYU：Normal , LONG：Normal |
| Intestinal organoid #4 | Human ESCs | RYU05 | Success | LONG05 | Success | RYU：Normal , LONG：Normal |
| Intestinal organoid #5 | Human iPSCs | RYU06 | Failure | LONG06 | Success | RYU：Proliferation of non-intestinal cells , LONG：Normal |
| Intestinal organoid #6 | Human iPSCs | RYU07 | Success | LONG07 | Success | RYU：Normal , LONG：Normal |
| Intestinal organoid #7 | Human iPSCs | RYU08 | Failure | LONG08 | Success | RYU：Non-proliferative , LONG：Normal |

**Figure S4. Generation of intestinal epithelial and mesenchymal stromal cells from intestinal organoids**
