## Supplementary material for "Human intestinal organoid-derived PDGFRα+ mesenchymal stroma empowers LGR4+ epithelial stem cells": Figure S5

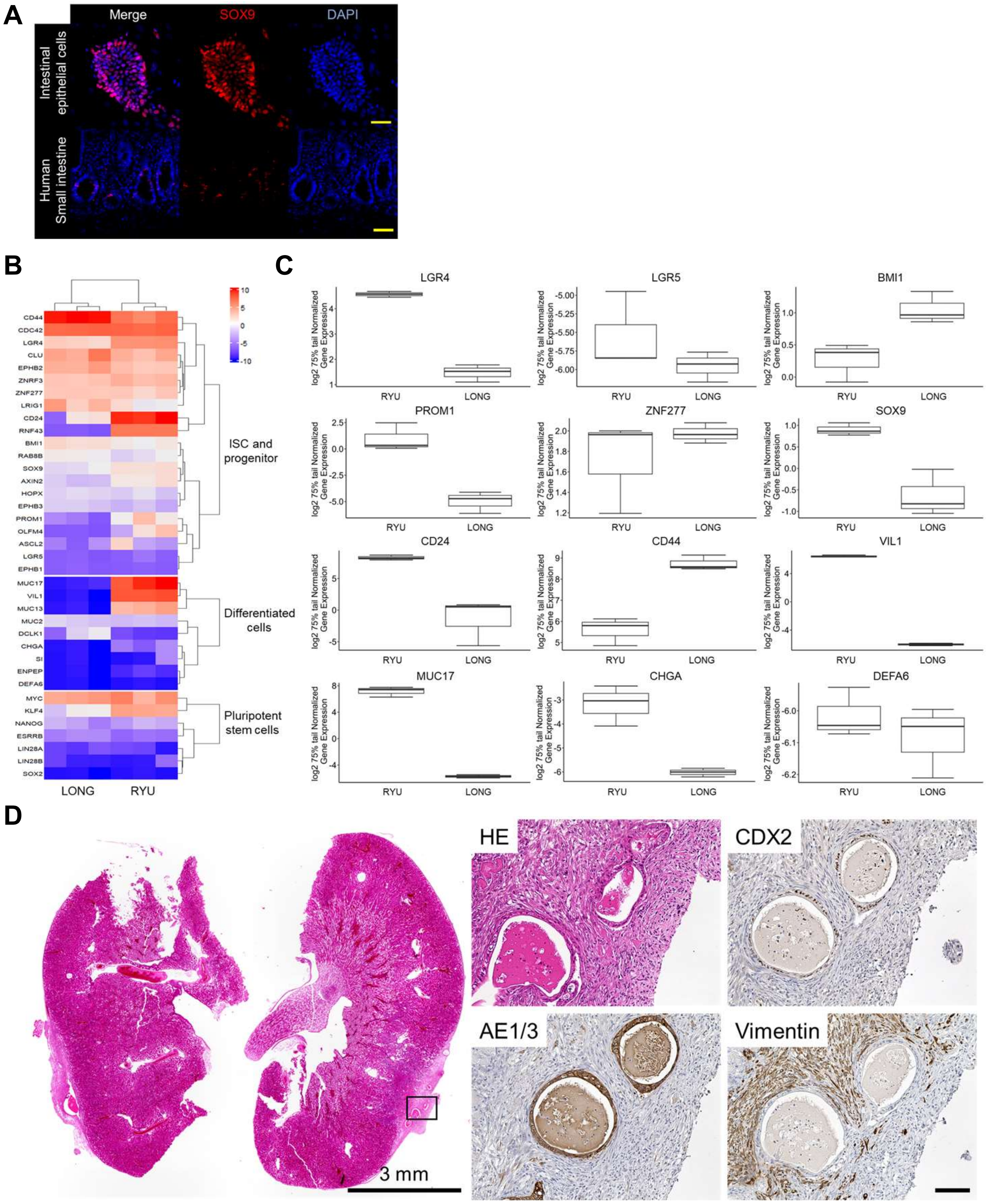


**Figure S5. Intestinal epithelial cells are positive for crypt base cell (CBC) markers such as LGR4, CD24, and CD44**
